# Cell-specific Na^+^ accumulation is linked to symplastic transport in tomato leaves

**DOI:** 10.64898/2026.03.26.714552

**Authors:** Lidor Shaar-Moshe, Daniel E. Runcie, Siobhán M. Brady

**Affiliations:** Department of Plant Biology and Genome Center, University of California, Davis, Davis, CA 95616 USA; Department of Evolutionary and Environmental Biology, Faculty of Natural Sciences, Institute of Evolution, University of Haifa, Haifa, Israel; Department of Plant Sciences, University of California, Davis, Davis, CA 95616, USA; Howard Hughes Medical Institute, UC Davis, Davis CA 95616 USA

**Keywords:** Na^+^ accumulation, symplastic transport, *plasmodesmata-located protein 1* (PDLP1), Solanum species, tomato, introgression line population, leaf

## Abstract

Soil salinization is a growing global threat that limits crop productivity. To cope with sodium (Na⁺) stress, plants have evolved tolerance mechanisms, including excluding Na⁺ from shoot tissues and tolerating elevated Na⁺ within shoots through tissue- and cellular-level mechanisms. Most current knowledge of Na⁺ accumulation comes from organ- or whole-plant measurements that lack the spatial resolution needed to resolve cellular tolerance mechanisms. Here, we used histological approaches to map leaf Na⁺ distribution in tomato (*Solanum*) species with contrasting salt-tolerance strategies. In the Na⁺-excluding domesticated tomato (cv. M82), Na⁺ was largely confined to the bundle sheath, whereas Na⁺-including wild relatives accumulated Na⁺ throughout the blade mesophyll. Consistent with these cell population-specific Na⁺ patterns, M82, but not *S. pennellii*, exhibited reduced symplastic transport and plasmodesmal permeability under salt stress. A genetic screen combined with transcriptome profiling implicated *Plasmodesmata-Located Protein 1* (*PDLP1*), a regulator of callose-mediated plasmodesmal closure, in establishing symplastic domains in M82 that restrict Na⁺ movement into the mesophyll. Moreover, *PDLP1* expression negatively correlated with mesophyll Na^+^ levels across wild and domesticated tomatoes. Collectively, these results link cellular Na⁺ enrichment patterns to symplastic connectivity and suggest that *PDLP1*-mediated regulation of plasmodesmata contributes to leaf-level salt-tolerance strategies.

**Highlights:**

1. Cell type-specific Na⁺ accumulation differs between domesticated tomato (*Solanum lycopersicum* cv. M82) and its wild relative *S. pennellii*.
2. Additional salt-tolerant wild tomato relatives exhibit leaf Na⁺ enrichment patterns similar to *S. pennellii*.
3. Salt stress reduces symplastic transport and plasmodesmal permeability in M82 leaves but not in *S. pennellii*.
4. An introgression line (IL6-4) between the two tomato species, which carries *S. pennellii Plasmodesmata-Located Protein 1* (*SpPDLP1*), shows *S. pennellii*-like Na⁺ enrichment patterns.
5. *PDLP1* expression shows a negative correlation with mesophyll Na^+^ levels across tomato species.

## Introduction

Salt stress is a growing environmental constraint on crop productivity, particularly in coastal, semi-arid, and arid regions. Climate change exacerbates soil salinization, which limits plant growth, especially in new shoots, by reducing the plant’s capacity to take up water. Additionally, excessive accumulation of sodium ions (Na^+^) within plants is toxic to key biochemical processes and accelerates the senescence of mature leaves (Munns & Tester, 2008; van Zelm et al., 2020). Approximately 33% of irrigated agricultural land is already salt-affected, and this area is expanding by ∼1-2 million hectares per year, in part due to the increasing use of treated wastewater and marginal-quality water for irrigation (Hopmans et al., 2021; Jenks et al., 2007). Thus, understanding and optimizing the molecular and cellular mechanisms underlying salt stress tolerance is critical for breeding crops with sustained productivity on salt-affected agricultural lands.

Upon entering the root epidermis through nonselective cation channels (transcellular transport) (Demidchik & Maathuis, 2007) Na^+^ can move into inner root cell layers via plasmodesmata (microchannels connecting the cytoplasm of adjacent cells; symplastic transport), diffuse through extracellular cell wall spaces (apoplastic transport), or be transported cell-to-cell via repeated membrane crossings (channels or transporters; transcellular transport) (van Zelm et al., 2020). To maintain cellular function and prevent damage to membranes and proteins (Benito et al., 2014), plants regulate cytosolic Na^+^ levels by exporting Na^+^ from root cells or sequestering it in root cell vacuoles through transporters and antiporters that sustain Na^+^/K^+^ balance (Almeida et al., 2017). Recently, Ramakrishna et al. (Ramakrishna et al., 2025) demonstrated that the highly selective sodium/proton antiporter SALT OVERLY SENSITIVE 1 is crucial for vacuolar Na^+^ sequestration, in addition to its role in Na^+^ extrusion into the cell wall. Once Na^+^ enters the root xylem, it travels with the transpiration stream through the stem and petiole xylem, potentially accumulating in the leaf blade over time. Since only a small portion of Na^+^ is recycled from shoot to root via the phloem, most Na^+^ reaching the shoot remains there, making the leaf blade the main site of Na^+^ toxicity for most plants (Munns & Tester, 2008). Na^+^ that accumulates in the blade mesophyll must first cross the bundle sheath, a parenchymatous cell layer surrounding the leaf’s vascular bundles, either symplastically or apoplastically. In some grass species, the bundle sheath cell walls are suberized and have been shown to restrict the movement of apoplastic dyes (Botha et al., 1982). However, another study using different apoplastic dyes challenged this finding (Mertz & Brutnell, 2014), raising questions about the role of the bundle sheath as an apoplastic barrier.

Given the detrimental effects of increased Na^+^ accumulation on photosynthesis and the potential to enhance crop growth and quality by manipulating leaf Na^+^ accumulation, it is surprising how little we know about the cellular domains of Na^+^ accumulation in leaves. A possible explanation for this knowledge gap is that salt-stress studies have largely overlooked the functional heterogeneity of distinct cell populations that comprise the above-ground plant body. Additionally, analyses of Na^+^ levels are traditionally conducted on bulk tissues using destructive (Hansen et al., 2013) or indirect methods (Arino-Estrada et al., 2019; Perelman et al., 2020), which either eliminate our ability or lack adequate resolution to identify the differential accumulation of Na^+^ in distinct cell populations and the underlying genetic mechanism(s). Thus, bridging these knowledge gaps requires elucidating the functions of distinct cell populations, the cellular patterning of Na^+^ accumulation, and the integration of these processes across the cellular and tissue levels to facilitate whole-plant salt tolerance.

Tomato is among the world’s most produced vegetable crops (Ritchie et al., 2023), cultivated in many arid and semi-arid regions, where brackish or saline water is used for irrigation. Under such irrigation, tomato growth and yield decline as water salinity increases (Zhai et al., 2015). Yet, their wild relatives are native to arid regions with high soil salinization and thus adapted to better withstand these unfavorable growth conditions (Bonarota et al., 2022). It has long been known that at the organ level, the domesticated tomato species attempts to exclude salt from its leaves, while wild tomatoes, including *S. pennellii*, accumulate salt in their leaves (Tal & Shannon, 1983). To dissect the phenotypic diversity and identify genetic loci regulating salt stress tolerance, researchers have utilized an introgression line population generated by crossing *S. pennellii* LA0716 with the processing inbred cv. M82 (Eshed & Zamir, 1994; Eshed & Zamir, 1995). This publicly available population has been genotyped at ultrahigh density to determine the exact gene content harbored within each line (Chitwood et al., 2013). A wide range of biologically and agronomically relevant traits, including salt stress tolerance (Frary et al., 2010; Frary et al., 2011; Li et al., 2011; Uozumi et al., 2012), have been previously analyzed using this IL population. However, the cellular and genetic basis for the divergent salt tolerance strategies of *S. pennellii* and *S. lycopersicum* cv. M82 (hereafter referred to as M82) is poorly understood.

In this study, we sought to investigate whether the divergent Na^+^ accumulation observed at the tissue level between the two tomato species is also evident at the cellular level. To achieve this, we employed a histological approach to identify domains of cellular Na^+^ accumulation in tomato leaves, revealing Na^+^ enrichment in distinct cell populations. While Na^+^ accumulation is enriched in the bundle sheath of M82, *S. pennellii* primarily accumulates Na^+^ in the blade mesophyll. These accumulation regions coincide with decreased symplastic transport and plasmodesmal permeability in salt-stressed M82 leaves, but not in *S. pennellii*. To elucidate the underlying genetic regulation, we screened the introgression line (IL) population derived from *S. pennellii* × M82 for salt tolerance and identified a single line, IL6-4, in which Na^+^ accumulation extended into the mesophyll, as observed in *S. pennellii*. Based on transcriptomic profiling (bulk RNA-seq) and real-time quantitative PCR (RT-qPCR), we found that *Plasmodesmata-Located Protein 1* (*PDLP1*), a negative regulator of symplastic transport (Thomas et al., 2008) is expressed at lower levels in IL6-4, and *S. pennellii* than in M82. Finally, we observed lower levels of *PDLP1* expression in two other salt-tolerant wild relatives of tomato, alongside an increased pattern of Na^+^ accumulation in the blade mesophyll. Our findings describe, for the first time in *Solanum* species, the preferential accumulation of Na^+^ in specific leaf cell populations. Furthermore, our work links divergent Na^+^ accumulation domains across tomato species to differential symplastic transport, providing a mechanistic model for foliar regulation of Na^+^ transport and accumulation at the cellular level.

## Results

### M82 and *S. pennellii* exhibit divergent Na^+^ accumulation patterns in their leaves

*S. lycopersicum* cultivars are generally moderately tolerant to salt stress, show reduced growth under such conditions, and use a Na^+^ exclusion mechanism to prevent Na^+^ buildup in their shoots. In contrast, their salt-tolerant wild relatives, including *S. pennellii,* exhibit less growth reduction under salt stress and typically employ a Na^+^ inclusion mechanism in their shoots (Dehan & Tal, 1978; Shannon et al., 1987). We first tested whether the two tomato species display different salt tolerance strategies (i.e., excluder vs. includer) under our experimental conditions. We exposed the plants to salt stress, starting at the early vegetative stage with three to four fully expanded leaves, and gradually increased the salt concentration in the nutrient-rich water by irrigating daily with 25, 50, and 75 mM NaCl until reaching a target concentration of 100 mM NaCl. This concentration was maintained for nine days, during which plants developed seven to nine leaves. Overall, the five-week-old plants were exposed to salt stress for two weeks. During the salt treatment, control plants were irrigated daily with nutrient-rich water (electrical conductivity of 1.4 dS/cm) and developed the same number of leaves as the treated plants. Consistent with previous reports, after two weeks of salt stress conditions, M82 shows a significant decrease in total biomass and leaf relative water content, along with an increase in osmotic adjustment (**Supplementary Fig. S1**). This adaptation is likely achieved through the synthesis of compatible solutes, as K^+^ levels moderately decrease and Na^+^ levels remain relatively low compared with *S. pennellii* (**Supplementary Fig. S2A&B**). In contrast to M82, *S. pennellii* maintained its biomass under salt stress and showed a minor increase in osmotic adjustment (**Supplementary Fig. S1**), likely due to the high Na^+^ levels accumulated in its shoots, as K^+^ levels sharply decreased under salt stress (**Supplementary Fig. S2A&B**). Together, these data show that M82 behaves as a shoot Na⁺ excluder that restricts Na⁺ entry into the leaves and largely preserves leaf K⁺, whereas *S. pennellii* acts as a shoot Na⁺ includer, accumulating high Na⁺ and allowing a lower K⁺/Na⁺ ratio in its leaves under salt stress (**Supplementary Fig. S2C&C’**) (Albaladejo et al., 2017; Tahal et al., 2000).

We hypothesized that these divergent salt tolerance strategies result from variations in cellular regulation of Na^+^ transport and accumulation between the two Solanum species. To test the hypothesis that cellular Na⁺ accumulation differs between species, we used CoroNa Green, a Na⁺-specific fluorescent dye (Shao et al., 2020). Because CoroNa Green fluorescence and membrane impermeability depend on living, intact cells, it was essential to ensure that cells in the sections remained viable during staining and imaging (**Supplementary Fig. S3A&B**). Using this Na⁺-specific dye, we found that, in M82 leaves exposed to two weeks of salt stress, Na⁺ predominantly accumulates in the bundle sheath, identified by the presence of amyloplasts, and in adjacent parenchyma cells on the abaxial side of the leaf (**Fig. 1A, Supplementary Fig. S3C-F**). The enhanced Na⁺ accumulation in the abaxial bundle sheath is consistent with early reports that, in tomato, the bundle sheath surrounds the veins only on the abaxial side (Venning, 1949). Moreover, this pattern supports growing evidence for differential ion accumulation and cell population-specific gene expression and function, implicating the bundle sheath as a regulatory, selective barrier that can restrict ion transport (Aubry et al., 2014; Shapira et al., 2009; Wigoda et al., 2014). In contrast, in *S. pennellii*, Na^+^ mainly accumulates in the mesophyll, particularly under salt stress. (**Fig. 1B&C**). Together, these results reveal species- and cell type-specific Na⁺ accumulation patterns that correspond with the different salt tolerance strategies of the two species (**Fig. 1A-C, Supplementary Fig. S1&2**).

**Figure 1.**
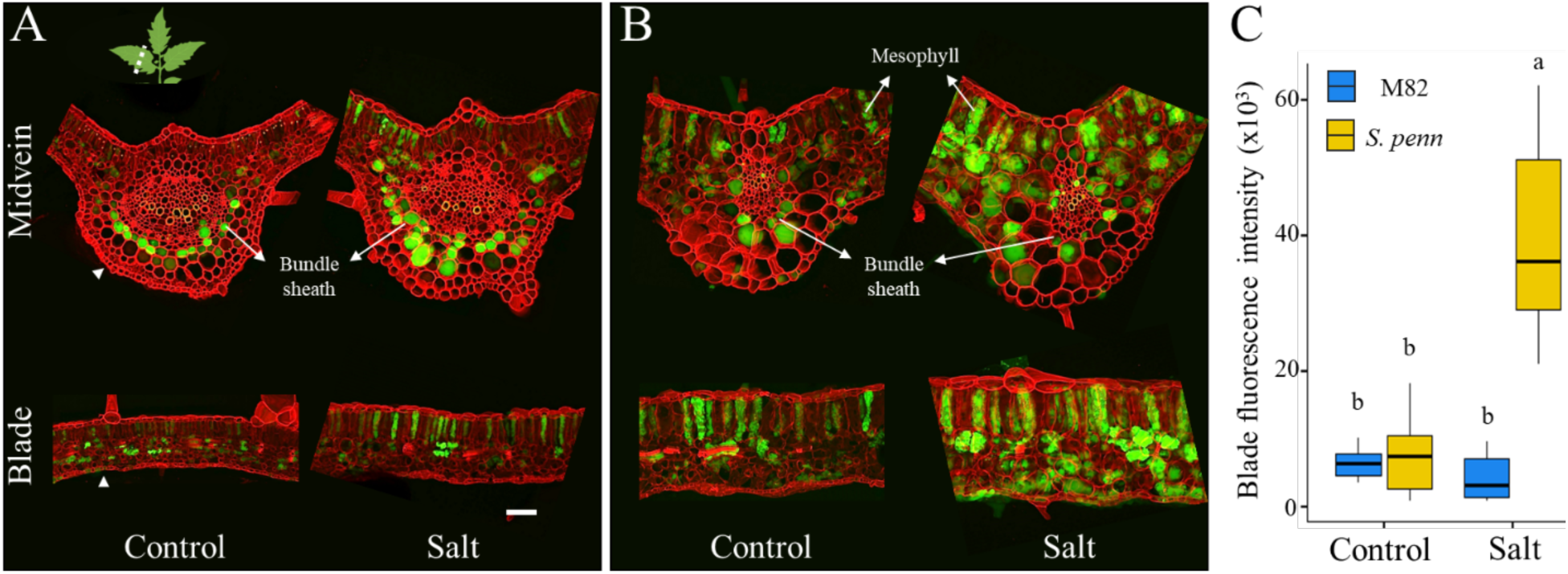
Wild and domesticated tomatoes accumulate Na^+^ in divergent leaf cell domains. Representative live cross-sections of the midvein and leaf blade from the third fully expanded leaf of five-week-old plants, stained with the Na⁺-specific dye CoroNa Green (green) and Congo Red (red) to visualize cell walls. Na^+^ preferentially accumulates (**A**) in M82 in bundle sheath and adjacent parenchyma cells at the abaxial side of the leaf. (**B**) In *S. pennellii*, Na^+^ accumulation is enriched in the blade mesophyll. Three-week-old plants were exposed to salt stress for two weeks, starting with 25 mM NaCl and increasing by 25 mM daily until reaching 100 mM NaCl, which was maintained for nine days. Scale bar indicates 100 µm (**C**) Quantification of CoroNa Green signal in blade mesophyll cells (n=6). Bars are median ± SD. Significant differences (p<0.05) within each condition, detected by two-way ANOVA followed by Tukey’s HSD test, are indicated by different letters.

### Enriched Na^+^ accumulation domains are conserved within each species and across other salt-tolerant wild relatives of tomato

Since Na^+^ accumulation is dynamic and occurs preferentially in older than younger leaves (Munns & Tester, 2008), we examined whether the observed cellular Na^+^ accumulation patterns are conserved within each species. To this end, we quantified the number of Na^+^-accumulating cells in six leaf cell populations from young and mature leaves (i.e., the primary leaflet of the youngest fully expanded leaf and its immediate predecessor), after five and nine days of salt stress, as well as in the corresponding control plants. Our analysis reveals that the enriched Na⁺-accumulation domains in the M82 bundle sheath and the *S. pennellii* mesophyll are characteristic of each species and are independent of leaf ontogeny (young or mature leaf), growth conditions (salt stress or control), and the severity of salt stress (five or nine days of exposure) (Fisher’s exact test, P < 0.01) (**Fig. 2A**). Given that *S. lycopersicum* has several wild relatives native to saline environments (Bonarota et al., 2022), we asked if the enriched Na^+^ accumulation in the blade mesophyll is unique to *S. pennellii* or conserved among additional salt-tolerant wild relatives of tomato. To this end, we selected two wild relatives of tomato: *S. pimpinellifolium* (LA1589), which belongs to the same clade as *S. lycopersicum*, and *S. peruvianum* (LA1954), which belongs to a separate clade (**Fig. 2B**). The three wild tomato species produced comparable biomass under control and salt conditions, as opposed to M82, which exhibited a significant biomass reduction under salt stress conditions (**Supplementary Fig. S4**). We quantified Na^+^ levels using CoroNa Green staining and found a significant Na^+^ accumulation in the blade mesophyll of *S. peruvianum*, with a similar pattern in *S. pimpinellifolium* under salt stress conditions (**Fig. 2C&D**).

**Figure 2.**
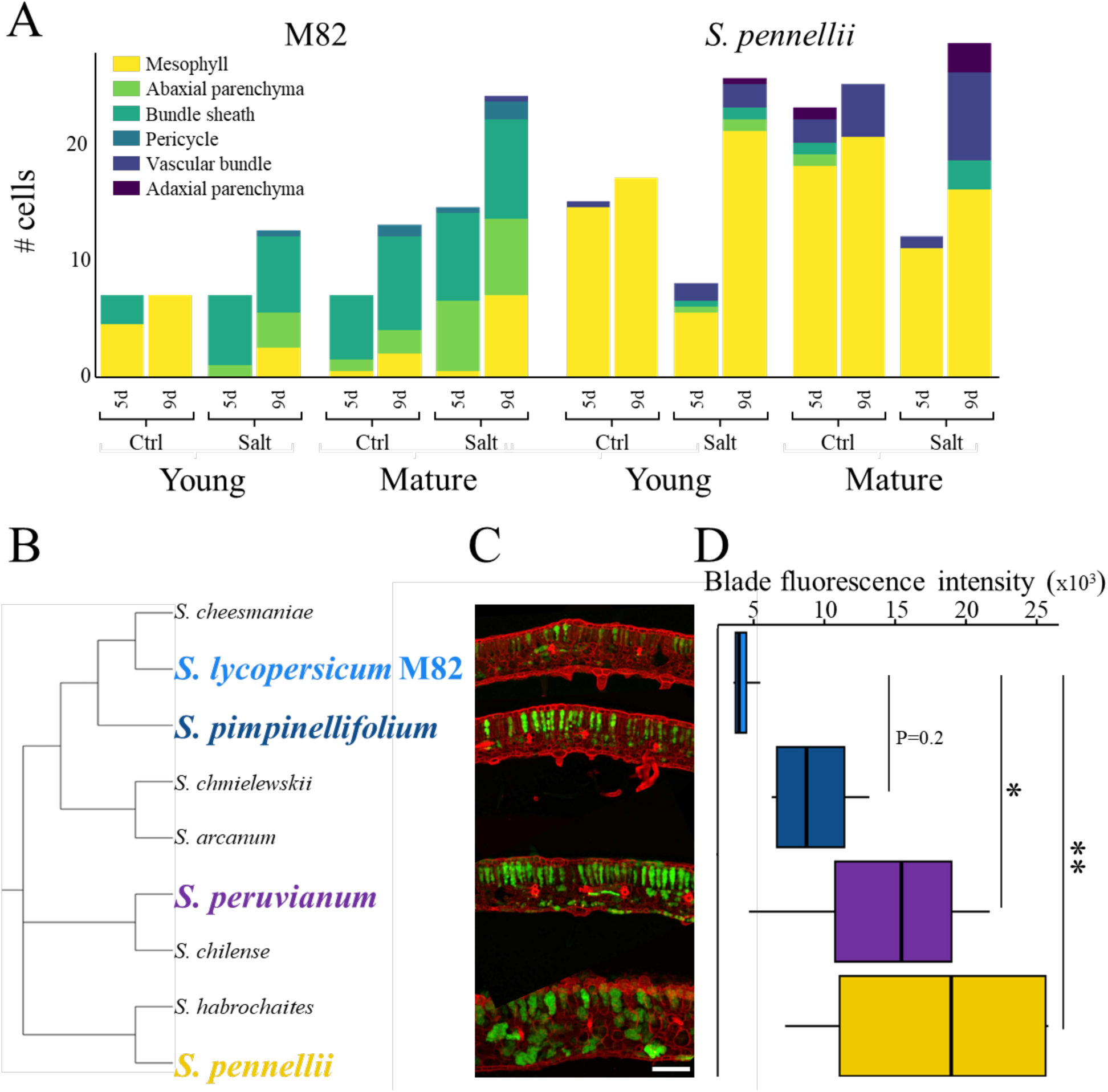
Leaf Na^+^ accumulation domains are conserved both within and across tomato species. (**A**) Stacked bar plots of the number of cells accumulating Na^+^ among different cell populations in live leaf cross sections, focused on the midveins of young and mature leaves, following five and nine days of salt stress and at the same time points under control conditions (n=6). Enriched Na^+^ accumulation domains within each species are conserved independent of the developmental stage of the leaf, growth conditions (salt stress or control conditions), or the duration of the salt stress. (**B**) Phylogenetic tree based on (Grandillo et al., 2011) of domesticated tomato (M82) and selected salt-tolerant wild relatives that are indicated in bold and colored based on the different tree branches. (**C**) Representative cross sections of live leaf blades from salt-stressed plants. Sections are stained with CoroNa Green (green), a Na^+^-specific dye, and Congo red (red), which stains the cell wall. The scale bar indicates 100 µm. (**D**) Quantification of CoroNa Green signal in blade mesophyll cells (n≥4). Significant differences detected by one-way ANOVA followed by Dunnett’s test are denoted with asterisks (** P≤0.05, and *** P≤0.01).

### Symplastic transport drives the differential Na^+^ accumulation domains

After Na⁺ is delivered to the leaf in the transpiration stream, it initially resides in the xylem apoplast, from where it may either remain within the apoplastic continuum surrounding the veins or be transported across the plasma membranes of xylem parenchyma and bundle sheath cells via Na⁺-permeable transporters and channels, enabling subsequent symplastic redistribution to the blade mesophyll. We hypothesize that the relative contributions of symplastic uptake, apoplastic retention, or apoplastic barriers vary between the two species and account for their distinct Na⁺ distribution patterns in the leaf. We therefore traced the symplastic and apoplastic pathways, along with the presence of consistent or inducible apoplastic barriers around their midveins. To test the hypothesis that symplastic transport regulates leaf Na^+^ accumulation patterns, we used the fluorescent symplastic tracer 5(6)-Carboxyfluorescein diacetate (CFDA) (Jiang et al., 2019) in dark-adapted tomato leaves, thus minimizing species differences in stomatal conductance and transpiration rates under salt-stress conditions (**Supplementary Fig. S5**). We observed a significant decrease in symplastic transport in M82 leaves under salt stress conditions, but not in *S. pennellii* leaves (**Fig. 3A, Supplementary Fig. S6A**). Since reduced symplastic trafficking results from decreased plasmodesmal permeability, we next used the drop-and-see (DANS) assay (Cui et al., 2015) to quantitatively assess whether the two species also exhibit differences in cell-to-cell permeability. Consistent with the previous assay, the spread of the fluorescent signal, which is proportional to plasmodesmal permeability, significantly decreased in leaves of salt-stressed M82 but not *S. pennellii* (**Fig. 3B, Supplementary Fig. S6B**). Lastly, we examined apoplastic transport using the apoplastic tracer berberine, which was chased with potassium thiocyanate to remove non-specifically bound berberine and immobilize it (Enstone & Peterson, 1992). This procedure allows us to qualitatively assess the functional apoplastic pathway in dark-adapted tomato leaves (**Supplementary Fig. S5**). Tracing the apoplastic transport showed that in M82 leaves, the dye was not consistently confined to the veins but accumulated in the mesophyll, where Na^+^ did not accumulate. In contrast, in *S. pennellii*, the apoplastic tracer accumulated in and around the veins, while Na^+^ accumulated in the mesophyll (**Fig. 3C**). Furthermore, staining with basic fuchsin or Auramine O, which binds hydrophobic cell wall components such as lignin and suberin that can accumulate in bundle sheath cells and restrict the apoplastic pathway (Botha et al., 1982), revealed no evidence of an apoplastic barrier around the midveins in either tomato species that can explain the patterns of Na^+^ accumulation (**Fig. 4D**, **Supplementary Fig. S8C**). Together, these results best support symplastic transport as the predominant driver of leaf Na⁺ patterning under our conditions.

**Figure 3.**
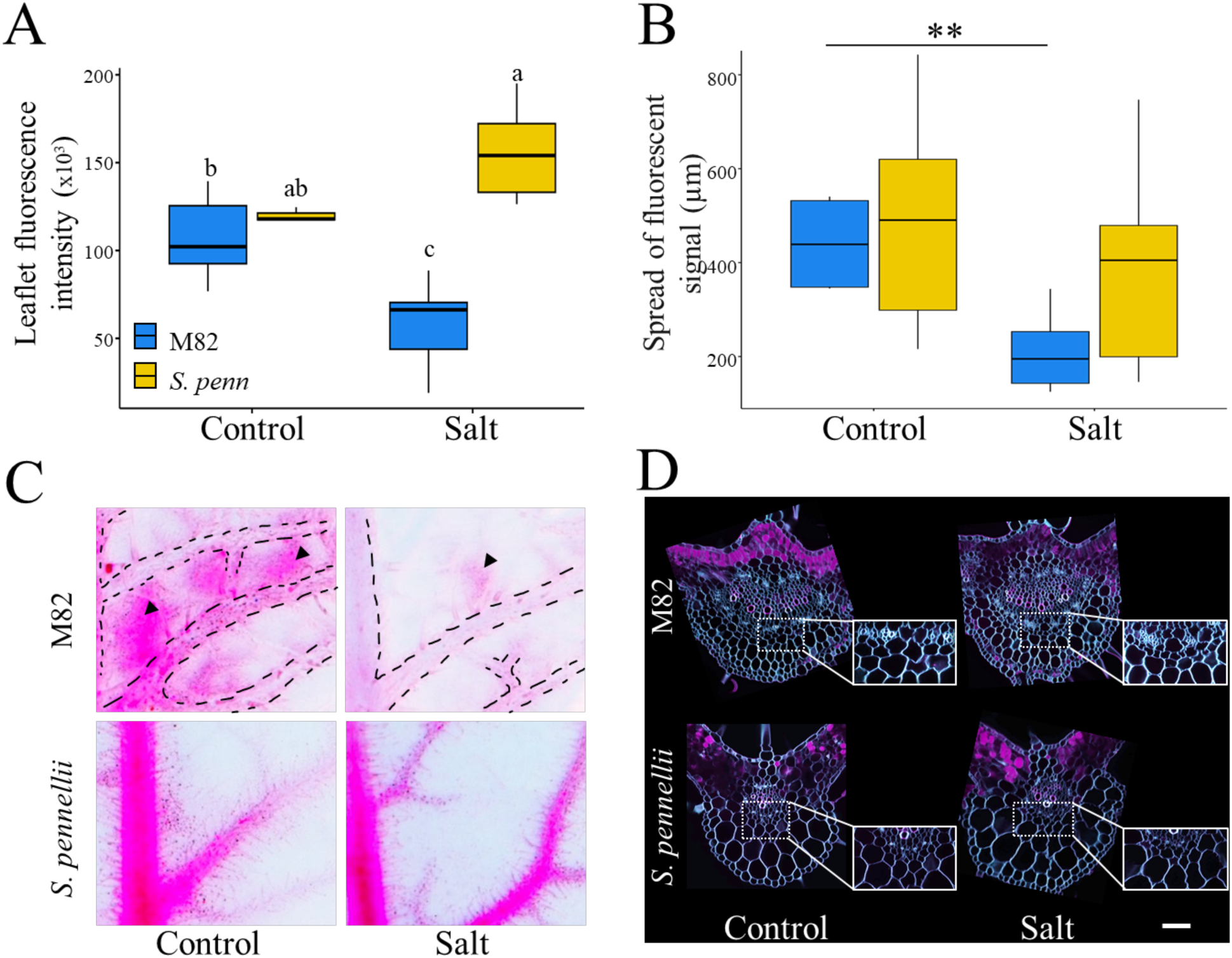
Symplastic, not apoplastic, transport primarily drives the different Na^+^ accumulation domains. (**A**) Quantification of the symplastic tracer 5(6)-Carboxyfluorescein diacetate (CFDA). Terminal leaflets of the youngest fully expanded leaves were subjected to 2h of dark acclimation following 2h of petiole feeding of CFDA (n=6). Bars are median ± SD. Significant differences (P<0.05), detected by two-way ANOVA followed by Tukey’s HSD test, are indicated by different letters. (**B**) Quantification of plasmodesma permeability based on CFDA diffusion. Significant differences (P<0.01), detected by Student’s t-test, are indicated by asterisks; ns indicates a non-significant difference. (**C**) Representative primary leaflets of the youngest fully expanded leaves were stained with the apoplastic tracer Berberine, followed by chasing with potassium thiocyanate for Berberine immobilization. Dashed lines indicate veins, and arrowheads indicate the tracer accumulation outside the veins. (**D**) Representative leaf cross sections of the midvein stained for lignin with Basic Fuchsin (magenta) and cellulose with Direct yellow (cyan). Autofluorescence of the chlorophyll is observed in magenta in mesophyll cells. Insets are high-magnification images of the boxed regions, focusing on the bundle sheath and adjacent parenchyma cells that lack consistent or inducible lignin deposition. The scale bar indicates 100 µm.

**Figure 4:**
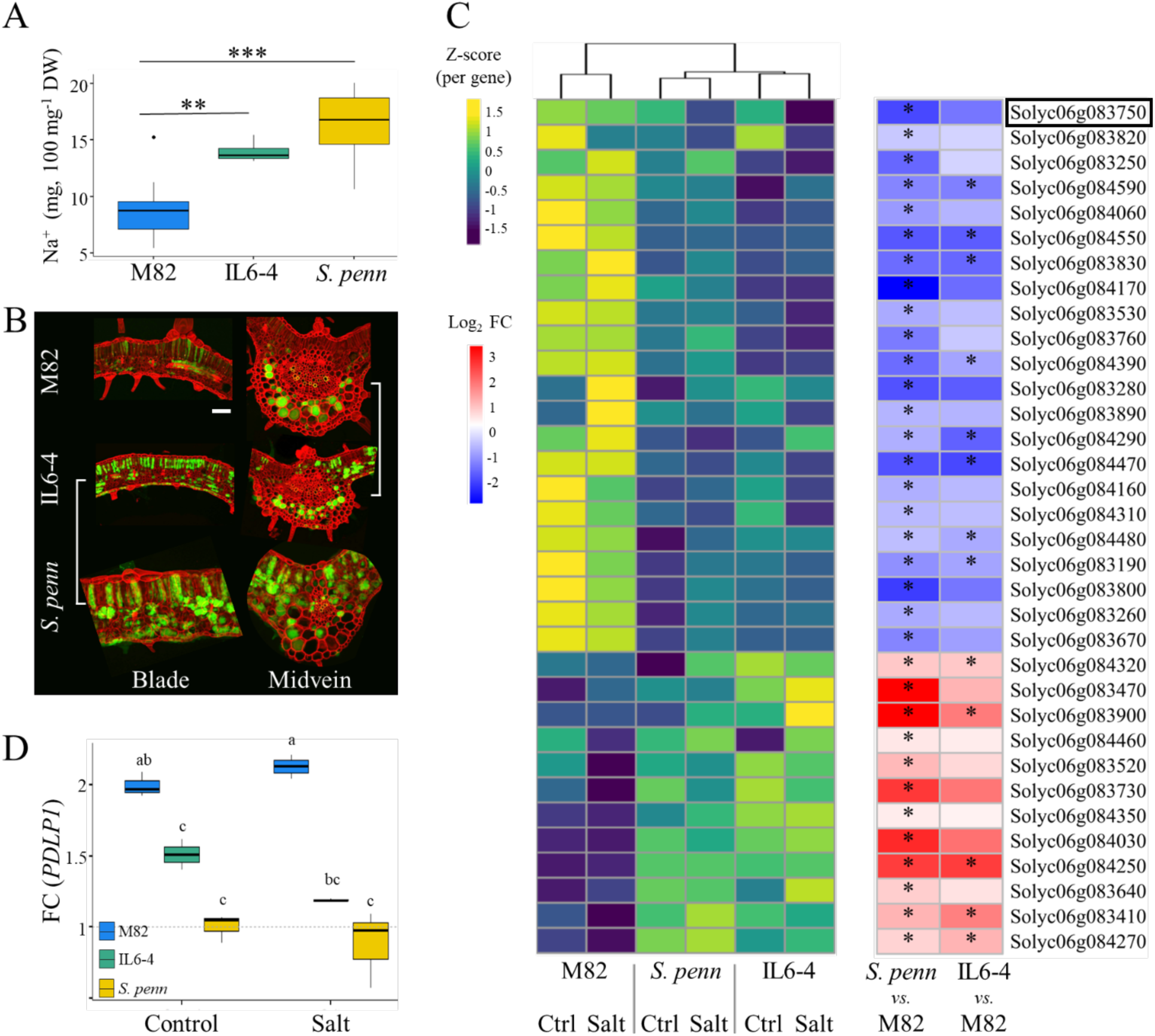
The Na^+^ accumulating introgression line 6-4 (IL6-4) harbors *SpPDLP1,* which shows reduced expression in IL6-4 and *S. pennellii* compared with M82. (**A**) Box plots of leaf Na^+^ content in plants subjected to salt stress conditions (n=6). Quantification was done with ICP-MS. Significant differences detected by one-way ANOVA followed by Dunnett’s test are denoted with asterisks (** P≤0.01, and *** P≤0.001). (**B**) Representative live cross sections of tomato midveins and leaf blades stained with CoroNa Green (green), a Na^+^ specific dye, and Congo red (red) that stains the cell wall. IL6-4 accumulates Na^+^ at the abaxial side of the midvein as the domesticated parent (M82) and in the mesophyll, similar to the wild parents (*S. pennellii*). Scale bar indicates 100 µm. (**C**) Heatmaps of expression levels (left) and fold change (FC) values (right) of 34 genes that introgressed from the wild parent into IL6-4 and are *i*) differentially expressed (FDR≤0.1) between *S. pennellii* and M82 under salt stress conditions, and *ii*) maintain similar FC directionality between IL6-4 and M82 under salt stress conditions. Expression was evaluated by bulk RNA sequencing (n≥3). Asterisks indicate differentially expressed genes (FDR≤0.1) between *S. pennellii* or IL6-4 and M82 under salt stress conditions. (**D**) Box plots of relative expression of *Plasmodesmata-located protein 1* (*PDLP1*; Solyc06g083750, and Sopen06g035160). Expression was evaluated by RT-qPCR (n≥3), using primary leaflets from the second fully expanded leaf. Exp (SGN-U346908) was used as a housekeeping gene. Bars are median ± SD. Significant differences (P<0.05), detected by two-way ANOVA followed by Tukey’s HSD test, are indicated by different letters.

### A single introgression line encompasses enriched Na^+^ accumulation domains of both tomato species

To elucidate the genetic basis underlying the species-specific Na^+^ accumulation domains, we utilized an introgression line (IL) population between M82 and *S. pennellii* (LA0716) (Eshed & Zamir, 1994; Eshed & Zamir, 1995). The population includes 76 homozygous lines, each containing a segment from the wild species (*S. pennellii*) in the background of the domesticated tomato (M82). We first screened for ILs that exhibit phenotypic stability, defined as a non-significant or only marginal biomass reduction under salt stress, similar to *S. pennellii*, while maintaining M82-like biomass under control conditions (**Supplementary Fig. S1A**). We hypothesized that such phenotypically stable ILs, which perform like *S. pennellii* under salt stress, would also accumulate higher levels of Na⁺ in their leaves (**Supplementary Fig. S2A**), while retaining M82-like productivity under non-stress conditions. Seven ILs met both criteria (**Supplementary Fig. S7**) and were selected for quantifying leaf Na^+^ levels using both ICP-MS and CoroNa Green (**Supplementary Fig. S8A&C**). Based on ICP-MS measurements, among the seven ILs, only IL6-4 accumulated higher levels of Na^+^ but similar levels of K^+^ compared to M82 under salt stress conditions (**Fig. 4A, Supplementary Fig. S8A&B**). Additionally, we found that Na⁺ quantification by fluorescence has a strong and positive correlation with ICP-MS measurements (r=0.75, P=0.03, **Supplementary Fig. S8C**), further validating the accuracy of the Na^+^ dye-based measurement method.

Tracing and quantification of leaf Na⁺ accumulation in IL6-4 showed that, unlike M82, Na⁺ extends from the abaxial side of the midvein into the leaf mesophyll, similar to *S. pennellii* (**Fig. 4B; Supplementary Fig. S8D&E**). Because IL6-4 contains enriched Na⁺-accumulation domains derived from both the wild and domesticated parents (adjusted P=0.07; **Supplementary Fig. S8E**), we hypothesized that introgressed *S. pennellii* gene(s) underlie the elevated Na⁺ levels and its movement into the IL6-4 blade. To test this, we sequenced the transcriptomes of the three genotypes under both control and salt-stress conditions (n=4; **Supplementary Fig. S9A–D**), focusing on differentially expressed genes (DEGs) within the IL6-4 introgressed region (Chitwood et al., 2013) that display *S. pennellii*-like expression patterns in IL6-4 and differ from M82. Of the 139 introgressed *S. pennellii* genes with M82 orthologs in IL6-4 (**Supplementary Table S2**), we identified 34 DEGs (FDR≤0.1) between salt-stressed *S. pennellii* and M82 whose salt-induced fold-change patterns in IL6-4 mirror those in *S. pennellii* rather than in M82 (**Fig. 4C, Supplementary Table S3**). We examined the DEG set for candidates involved in ion transport and compartmentation, particularly via the symplastic pathway. Most DEGs were associated with general metabolism, RNA processing, or protein turnover (**Supplementary Table S3**). However, one gene, Solyc06g083750, encodes *PLASMODESMATA-LOCATED PROTEIN 1* (*PDLP1*; **Supplementary Fig. S9E**). In Arabidopsis and aspen, PDLP1 promotes callose deposition at plasmodesmata and thereby reduces plasmodesmatal permeability; elevated *PDLP1* levels are generally associated with restricted symplastic transport (Thomas et al., 2008; Tylewicz et al., 2018). Consistent with this role, *PDLP1* expression is higher in M82 leaves than in *S. pennellii* and IL6-4, as confirmed by real-time quantitative PCR (RT-qPCR) (**Fig. 4D; Supplementary Table S3**). This elevated *PDLP1* expression in M82 is in line with a more restricted symplastic pathway (**Fig. 3A&B**) and thus supports the confined Na⁺-accumulation domain at the abaxial side of the veins and the lower Na^+^ accumulation in M82 leaves (**Fig. 1A&C, Supplementary Figs. S2A, S8D&E**). Conversely, the lower *PDLP1* expression in *S. pennellii* and IL6-4 is consistent with enhanced symplastic connectivity in *S. pennellii* (**Fig. 3A&B**), which could facilitate Na⁺ movement from the bundle sheath into the mesophyll and thereby contribute to the more extended Na⁺-accumulation domains and overall higher leaf Na^+^ levels observed in these genotypes (**Figs. 1A&C, 3A&B, Supplementary Figs. S2A, S8D&E**).

Additional family members related to *PDPL1* in M82 and *S. pennellii*, as well as their Arabidopsis orthologs, were identified by phylogenetic analysis (**Supplementary Fig. S9E**). Among the eight *PDLP* members, two additional *PDLPs* (Solyc04g007950 and Solyc09g090680) exhibit induced expression under salt stress conditions in both M82 and IL6-4 compared with *S. pennellii* (FDR≤0.01, **Supplementary Fig. S9F, Table S1**), positioning these *PDLPs* as putative regulators of Na^+^ accumulation at the abaxial side of the midvein.

### *PDLP1* expression is negatively correlated with Na^+^ buildup in the blade mesophyll

Based on the known role of *PDLP1* in establishing symplastic domains (Bayer et al., 2008; Tylewicz et al., 2018), and our findings on divergent *PLDP1* expression, symplastic transport, and Na^+^ accumulation patterns between the two species, we propose that *PDLP1* is involved in the formation of leaf Na^+^ accumulation domains by regulating symplastic transport. We hypothesize that if increased *PDLP1* expression restricts leaf Na^+^ movement and reduces its accumulation in M82 mesophyll, then wild tomato relatives with higher Na^+^ accumulation in the blade mesophyll should exhibit lower *PDLP1* expression. Using RT-qPCR to analyze *PDLP1* orthologs, we observed lower and similar expression levels among the three wild tomato species compared to M82 under salt stress conditions (adjusted P≤0.066, **Fig. 3D**). Furthermore, the expression level of *PDLP1* orthologs is inversely related to Na^+^ accumulation across the four tomato species (r=-0.62, P=0.05, **Fig. 3E**). Collectively, these findings suggest that increased Na^+^ accumulation in the mesophyll of the tested salt-tolerant wild tomatoes is associated with lower *PDLP1* expression, indicating the absence of symplastic domains around the veins in these species.

## Discussion

Salinized soils are estimated to increase at an approximate rate of 10% per year, primarily due to agricultural practices (Hopmans et al., 2021). The expansion of soil salinity has rendered vast areas unusable for agriculture, reducing yields and shifting ecosystems toward halotolerant species, directly undermining food security (Vengosh, 2013). To mitigate yield decline and promote sustainable agriculture, plant researchers have attempted to increase crop salt tolerance by transferring genetic information from related salt-tolerant species. Tomatoes serve as a good example for such research efforts as they are primarily cultivated in arid and semi-arid areas, and have wild relatives endemic to arid coastal regions that can withstand high salt concentrations (e.g., *S. pimpinellifolium*, *S. pennellii*, *S. peruvianum*) (**Supplementary Figs. S1, S2&S4**) (Bonarota et al., 2022). Despite numerous QTL discoveries and the development of introgressed or transgenic plants, commercially successful salt-tolerant cultivars that include wild tomato alleles remain rare. This may stem from the polygenic and environment-dependent nature of the salt stress response. (Bonarota et al., 2022; Guo et al., 2022). Here, we propose that the divergent patterns of leaf Na^+^ accumulation between domesticated and wild tomato species (**Figs. 1&2**) may play a significant role in tissue tolerance and should be considered in breeding efforts to develop salt-tolerant crops varieties.

Several studies have demonstrated cell population-specific expression of Na^+^ transporters, which contribute to the directional retrieval of Na^+^ primarily from xylem vessels via xylem-parenchyma cells. Examples include *HKT1;1* from Arabidopsis, which is expressed in stelar root cells and in leaf parenchyma and is responsible for retrieving Na^+^ from xylem vessels (Sunarpi et al., 2005). A conserved expression pattern and effect on Na^+^ level in aerial tissues were reported for rice *HKT1;5,* and tomato *HKT1;2,* which are expressed in cells surrounding xylem vessels in roots and shoots, and contribute to plant salt tolerance (Jaime-Perez et al., 2017; Kobayashi et al., 2017). In all cases, organ-level Na^+^ accumulation was determined by destructive methods on bulk tissues, which averages out Na^+^ spatial heterogeneity, thereby impeding analysis of cellular functions and the underlying genetic regulation of salt tissue tolerance. A handful of studies have attempted to assess leaf ion accumulation at the cellular level using various techniques, including cryo-scanning electron microscopy (SEM), X-ray microanalysis, and sodium-specific dyes. In salt-treated barley (salt-tolerant cultivar) and wheat (salt-sensitive cultivar), Na⁺ accumulates in leaf tissues but shows no evidence of differential distribution between cell types. This contrasts with other ions, such as Cl⁻, which is preferentially accumulated in epidermal cells, and K⁺, which is more concentrated in mesophyll cells. (James et al., 2006). In contrast, in salt-treated *Puccinellia peisonis*, Na^+^ preferentially accumulates in the bundle sheath (Stelzer, 1981); in Rhodes grass (*Chloris gayana*), it accumulates in the xylem parenchyma and epidermis (Oi et al., 2022); and in citrus, Na^+^ mainly accumulates in the mesophyll (Gonzalez et al., 2012). These observations suggest that some plants accumulate Na^+^ in distinct leaf cell populations in a species-specific manner, supporting our findings (**Figs. 1&2**). Yet, further research employing standardized methodologies and additional species is needed to identify the cellular and molecular mechanisms underlying enriched Na^+^ accumulation domains in leaves. Recently, Ramakrishna et al. (Ramakrishna et al., 2025) demonstrated the use of a newly developed cryo-nanoscale secondary ion mass spectrometry (cryoNanoSIMS) ion microprobe to measure in parallel the subcellular localization of several ions in root meristems. Extending cryoNanoSIMS to leaves, together with a single-cell RNA approach, could help link cell population-specific transcriptional regulation to Na⁺ and other ion distributions.

Once exiting the xylem vessels in the leaf, xylem sap, carrying dissolved ions, can cross the cell layers surrounding the vascular bundle and progress from the bundle sheath toward the mesophyll and epidermis via the symplastic, apoplastic, and transcellular pathways. Several studies suggest that Na^+^ flow throughout the leaf is mainly symplastic and strictly regulated (Shapira et al., 2009; Speer & Kaiser, 1991) (**Figs. 1&3**). The regulation of symplastic connectivity depends on the permeability of plasmodesmata, which is influenced by callose deposition, an enzymatic process mediated by a family of callose synthases and associated regulatory proteins, including plasmodesmata-located proteins (PDLPs) (Thomas et al., 2008; Zavaliev et al., 2011). Such restriction of cell-to-cell trafficking establishes symplastic domains that are involved in fundamental developmental processes, including the formation of lateral roots under optimal conditions (Benitez-Alfonso et al., 2013) or when roots are not in contact with water (xerobranching, (Mehra et al., 2022), as well as photoperiodic control of bud dormancy (Tylewicz et al., 2018). Based on our results, we hypothesize that the restriction of symplastic trafficking in M82 is mediated by localized expression of *PDLP1* at midvein junctions with second-order veins, thereby limiting Na^+^ spread into the blade mesophyll and promoting its enrichment on the abaxial side of the midveins (**Fig. 5C**). The decreased plasmodesmal permeability in salt-treated leaves of M82 is accompanied by increased expression of *SlPDLP1*, in contrast to *S. pennellii* and IL6-4, which show lower expression levels of *SpPDLP1* and an extended Na^+^ accumulation domain in the photosynthetically active mesophyll cells of the blade. In accordance with the lower *SpPDLP1* expression and extended Na^+^ accumulation, there is no change in plasmodesmal permeability in salt-treated *S. pennellii* leaves (**Figs. 1,2&4**). Furthermore, our findings demonstrate an inverse relationship between the expression levels of *PDLP1* orthologs in two additional salt-tolerant wild relatives of tomato and the enrichment patterns of Na^+^ accumulation in their blade mesophyll (**Figs. 2 & 3**). Thus, we propose to expand the role of symplastic domains from developmental processes to tolerance strategies in response to salt stress.

**Figure 5.**
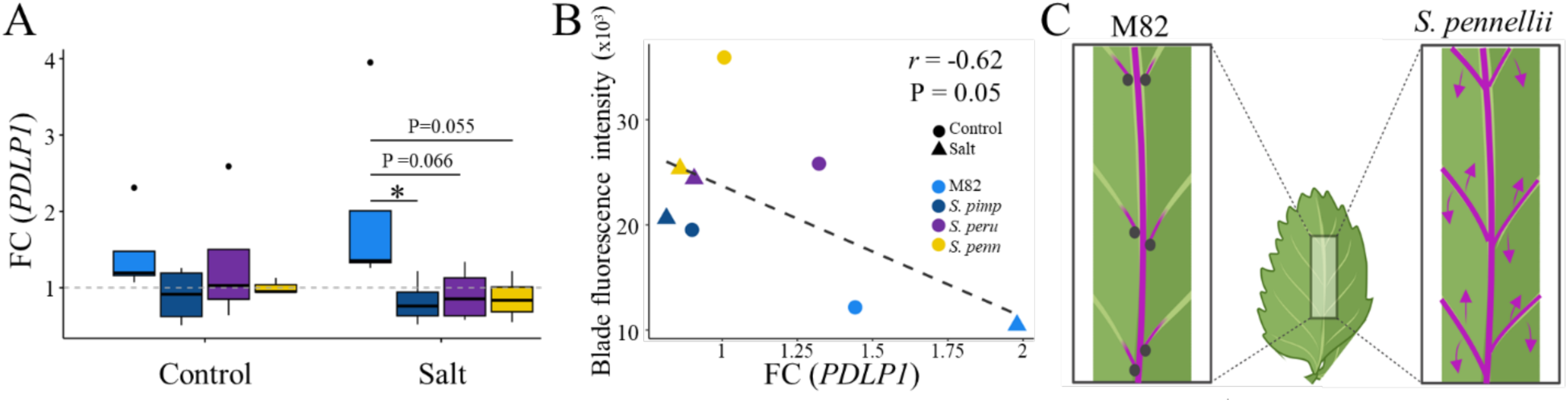
Additional salt-tolerant wild relatives of tomato with increased Na^+^ accumulation in the blade mesophyll also exhibit reduced expression of *PDLP1* orthologs. (**A**) Box plots of relative expression of *Plasmodesmata-located protein 1* (*PDLP1*) orthologs (n=4). Expression was evaluated by RT-qPCR (n≥3) using primary leaflets from the second fully expanded leaf. Exp (SGN-U346908) was used as a housekeeping gene. Bars are median ± SD. Significant differences detected by one-way ANOVA followed by Dunnett’s test are denoted with asterisks (* P≤0.05). (**B**) Spearman correlation between mesophyll Na^+^ accumulation (Fig. 2D) and *PDLP1* expression (**A**) across tomato species grown under control and salt stress conditions. (**C**) Schematic illustration of the working hypothesis. In M82, the increased and localized expression of *PDLP1* at midvein junctions with second-order veins establishes a symplastic domain that limits cell-to-cell transport of Na^+^ and facilitates its accumulation mainly in bundle sheath and adjacent parenchyma cells. In *S. pennellii*, low *PDLP1* expression enables symplastic trafficking and Na^+^ accumulation in mesophyll cells near the site of transpiration.

Given the paucity of high-resolution spatial information on Na^+^ accumulation in leaves, this work provides a comprehensive characterization of cellular Na^+^ accumulation patterns in wild and domesticated tomato species, which aligns with their well-documented differences in Na^+^ accumulation at the organ level and their divergent salt tolerance strategies at the whole plant level (i.e., Na^+^ includer vs. excluder) (Tal & Shannon, 1983). While *PDLP1* expression correlates with foliar Na⁺ patterning, establishing causality will require cell-type-specific perturbations of *PDLP1* and cell-resolved mapping of its expression domain. The proposed cellular mechanism provides a framework for a better understanding of the regulation of Na^+^ transport and accumulation, which is necessary to facilitate the precise engineering of tolerance mechanisms to enhance the performance of tomatoes and other vegetable crops under elevated soil salinity.

## Materials and methods

### Plant material

Seeds of *Solanum lycopersicum* cultivar M82, *S. pennellii* (LA0716), *S. pimpinellifolium* (LA1589), and *S. peruvianum* (LA1954) were obtained from the C.M. Rick Tomato Genetics Resource Center at UC Davis (https://tgrc.ucdavis.edu/). Seeds of the introgression line population between M82 and *S. pennellii* LA0716 were obtained from the Zamir lab (Eshed and Zamir, 1994, 1995).

### Salt stress experiments

Tomato seeds were surface-sterilized with 70% Ethanol for five minutes, followed by 20 minutes in a fresh 50% bleach solution, and rinsed five times with sterile water. Seeds were sown in 12×12 cm square Petri plates containing 4.3 g/L MS medium without vitamins (Caisson, MSP09-50LT), 1% (w/v) sucrose, 0.5 g/L MES (pH 5.8) and 1% (w/v) agar (Difco, 214530). Uniform, ∼6-9-day-old seedlings were transferred into 656 mL containers (D40L STUEWE) filled with Profile^®^ Field & Fairway™ (Turface Athletic Profile Field & Fairway clay substrate) and covered with a 2 cm layer of vermiculite. Plants were grown in a Conviron walk-in chamber (22°C, 70% RH, 16/8 hour light/dark cycle, light intensity of ∼150 µmol/m²/s) and irrigated every other day with nutrient-rich water (N:P:K, 2:1:2). A completely randomized design, with six replicates per treatment and species, was employed starting at three-four fully expended leaves (∼3-week-old seedlings). Plants were progressively exposed to salinity by irrigating daily with increasing salt concentrations, starting with 25 mM NaCl and increasing by 25 mM NaCl increments until reaching a target concentration of 100 mM NaCl, which was maintained for nine days. During this time, plants were watered daily with nutrient-rich water. Altogether, plants were exposed to salt for 14 days. Runoff electric conductivity was monitored and recorded daily once the target concentration was reached. During the salt treatment, control plants were irrigated daily with nutrient-rich water.

### Screening for Salt Tolerance Introgression Lines (ILs)

IL seeds were divided into five batches based on their corresponding chromosomes (i.e., chromosomes 3, 4, & 8; 2, 5, & 6; 10, 11, & 12; 7 & 9; 1 & low germinating ILs). Each screening assay included the ILs from each batch and the two parents: M82 and *S. pennellii* (LA0716). Seeds were surface sterilized with 50% HCl for 15 minutes, followed by 15 minutes in a 10% TPS solution, then rinsed and sown as described above. Uniform ∼6-9-day-old seedlings from each IL (n=20), M82 (n=40), and 32 *S. pennellii* (n=32) were transplanted into six 72-pot trays filled with profile mix (Turface Athletic Profile Field & Fairway clay substrate) in a completely randomized design, and watered with nutrient-rich water. At the stage of two fully expanded leaves, half of the plants were exposed to salt stress by irrigating with 50 mM NaCl. After three days, the trays were filled with 100 mM NaCl, which was replaced every two days. In total, plants were subjected to salt treatment for 12 days. During the salt treatment, control plants were irrigated daily with nutrient-rich water.

### Physiological Measurements

On the day of harvesting, Solanum species had seven to nine leaves. A fully expanded leaf is defined operationally as a leaf whose terminal and first pair of primary leaflets are completely spread. On harvest day, shoot and washed root samples (n = 6) were oven-dried (60°C for 4 days) and weighed to determine total plant biomass. The terminal leaflet from the third fully expanded leaf, located below the shoot apex, was collected for measuring relative water content as described in (Canto Pastor et al, 2024). Midday measurements of osmotic potential were made on a primary leaflet from the second fully expanded leaf. Leaflets were placed in vials containing deionized water, rehydrated in darkness at 4 °C for 6 hours to achieve full turgor, then blotted dry and snap-frozen in liquid N₂. Sap osmotic potential was determined with a vapor-pressure osmometer (Vapro 5600, Wescor/METER, USA); osmolality (cₒₛₘ, mOsm kg⁻¹) was converted to osmotic potential at full turgor as Ψₛ¹⁰⁰ (MPa) = −0.00248 × cₒₛₘ (25 °C). Osmotic adjustment (OA) was calculated as OA = Ψₛ¹⁰⁰ control - Ψₛ¹⁰⁰ stress, so that positive values indicate greater solute accumulation. Photosynthesis rate (A), stomatal conductance (gs), and transpiration rate were measured between 9:30 and 11:30 AM under light conditions and following 2 hours of dark adaptation on the abaxial surface of the terminal leaflet from the third or fourth fully expanded leaf using a LICOR-6400XT portable photosynthesis system (LI-COR Biosciences, Lincoln, Nebraska, USA). Light intensity was kept at 1,000 μmol m^−2^ s^−1^, with a constant air flow rate of 400 μmol s^−1^ and a reference CO_2_ concentration of 400 μmol CO_2_ mol^−1^ air.

### ICP-MS Elemental Analysis

On the day of harvesting, leaves and washed root samples of M82 and *S. pennellii* were oven-dried (60°C for 96 h), homogenized, and measured for their dry weight (DW). Samples were digested in concentrated nitric acid and analyzed by inductively coupled plasma mass spectrometry (ICP-MS) at the UC Davis/Interdisciplinary Center for Plasma Mass Spectrometry (UCD/ICPMS) to measure the relative mass of Na and K within each tissue. Leaf samples of M82, *S. pennellii,* and seven selected ILs from the salt tolerance screening assays were prepared the same way for ICP-MS, and measured at the UC Berkeley College of Natural Resources Inductively Coupled Plasma Spectroscopy Facility. Each genotype, tissue, and condition had at least six biological replicates.

### Histochemistry and Imaging

#### Sodium Staining and Viability Staining

Relative Na^+^ enrichment within foliar cell populations was determined following 12 or 14 days of salt treatment (see salt tolerance screening assays or salt experiments), using live leaflet cross-sections from the third fully expanded leaf within 24 to 32 hours of sampling. To assess the conservation of Na^+^ enrichment within distinct cell populations between M82 and *S. pennellii*, primary leaflet samples were collected from the first fully expanded leaf (i.e., young leaf) and third to fifth fully expanded leaves (i.e., mature leaves), following five, nine, and 14 days of exposing plants to salty water. In parallel, leaflets from control plants were sampled at the corresponding developmental stages.

One cm cuts from the middle third of the leaflet spanning the midvein were embedded in 4% (w/v) agar plugs (n=6). Plugs were kept in scintillation vials filled with fresh incubation buffer (20 mM MOPS [Thomas Scientific, C791M00], 0.5 mM calcium sulfate [Spectrum, CA171], and 200 mM sorbitol [Amresco, 0691]). Within two hours of sampling, 300 µm sections were prepared using a vibratome (Leica, VT1200S) and stored in a 12-well plate containing incubation buffer. Fifty micrograms of CoroNa Green were first dissolved in 100 μl dimethyl sulfoxide to prepare the stock solution. Four sections (i.e., technical replicates) from each genotype and treatment were transferred to a new well and incubated for ∼18 hours in the dark at room temperature, with gentle shaking, in 250 µL of 0.1 mM CoroNa Green reagent (ThermoFisher, C36676) diluted in incubation buffer. In parallel, three sections from each genotype and treatment were transferred to a new well containing incubation buffer alone and served as unstained controls for autofluorescence and viability tests. Before imaging Na^+^ accumulation or control sections to assess background autofluorescence, freshly prepared 0.15% (w/v) Congo red (MP, 105099), reconstituted in incubation buffer, was added to a slide containing the section and incubated for ∼1 minute. Congo red, which stains cellulose, was used to mark the different cell layers of the cross-section (Mitra & Dominique, 2014).

Samples were imaged with a Zeiss LSM 900 confocal microscope (water immersion, ×20 objective), with excitation at 488 nm and emission collection of 493-524 nm for CoroNa Green and 591-650nm for Congo red. Z-series sections were imaged at 4.5-μm intervals.

Because cellular sodium sequestration requires live cells, we validated cell viability of our sections using fluorescein diacetate (FDA; Sigma-Aldrich, F7378), whose fluorescence depends on cell membrane integrity (Jones et al., 2016). A fresh stock solution of 0.2% (w/v) FDA in acetone was prepared, diluted to 4 μg/mL in incubation buffer, and kept in the dark. Since the FDA signal quickly fades, samples were incubated for ∼30 minutes in the dark at room temperature in a fresh working solution of FDA and then imaged using a Zeiss LSM 700 confocal microscope (water immersion, ×20 objective), with excitation at 488 nm and emission detected using the eGFP filter at 500-550 nm. Z-series sections were imaged at 4.5-μm intervals. Autofluorescence signals from unstained samples were collected using identical imaging settings as those used for stained samples.

#### Leaf Cellular Anatomy Characterization

Characterization of tissue anatomy and Na^+^-accumulating cell types at the abaxial side of the midvein was performed on M82 leaflets from the third fully expanded leaf (n=6) using Toluidine Blue O (TBO, Baker, W143-03) and iodine-potassium iodide (IKI). Sections, either 300 or 150 µm for IKI and TBO, respectively, were fixed overnight with Formalin-Aceto-Alcohol (FAA) solution (v/v; 50% ethanol 95%, 5% glacial acetic acid, 10% formalin, 35% water) followed by gradual rehydration by 30 min incubations in 70%, 50%, 30% and 10% (v/v) ethanol. Fully rehydrated samples were stored in water at 4°C. To delineate leaf anatomy and visualize tomato’s bicollateral veins, we used 0.1% TBO (w/v), a metachromatic dye that is used to differentiate between primary and secondary cell walls and assist in orienting the position of the xylem, phloem, and surrounding cell layers (Yeung, 1998). Amyloplast-accumulating bundle sheath cells were detected using the starch-specific dye IKI, following the protocol specified in Yeung (Yeung, 1998). Sections were imaged using a bright-field Olympus AH-2 microscope (water immersion, ×20 objective).

Fixed sections (FAA, 150 µm) of M82 and *S. pennellii* primary leaflets from the third and fourth fully expanded leaf (n=6) were used to determine if cell walls of the bundle sheath and adjacent parenchyma contain hydrophobic compounds. To this end, we used basic fuchsin (Sigma, 857343-25G), together with Direct Yellow 96 (Biosynth, FD41192), which stains lignin and cellulose, respectively, and Auramine O (Sigma-Aldrich, 861030), a lipophilic fluorescent dye that stains suberin and other hydrophobic compounds, such as cutin. Following Ursache et al. (Ursache et al., 2018), for lignin and cellulose staining, sections were stained in Clearsee solution with 0.2% (w/v) basic fuchsin for 30 minutes, washed once with Clearsee solution, and then stained with 0.1% (w/v) Direct Yellow 96 (in Clearsee) for one hour. Sections were then washed twice with Clearsee solution and imaged with a Zeiss LSM700 confocal microscope (water-immersion ×20 objective). Basic fuchsin was imaged with 555 nm excitation, and emission collection of 600-650 nm. Direct Yellow 96 was imaged with 488 nm excitation, and emission collection of 500-550 nm. For suberin staining, sections were incubated in Clearsee solution containing 0.5% (w/v) Auramine O overnight, then washed twice for two hours in Clearsee solution, and finally imaged using a Zeiss LSM700 confocal microscope (water-immersion ×20 objective). Auramine O was imaged with 488 nm excitation, and emission collection of 505-530 nm. Autofluorescence was imaged with 555 nm excitation and emission collected between 560 and 800 nm.

#### Transport Assays and Plasmodesmal Permeability

Tracing of leaf apoplastic and symplastic transport pathways was done using primary leaflets from the second fully expanded leaf. Apoplastic tracing is based on Enstone and Peterson (Enstone & Peterson, 1992), with the following modifications: Following 14 days of salt exposure, M82 and *S. pennellii* plants (n=6) were transferred to a dark room for 2 hours to minimize differences in transpiration rates resulting from growth conditions. Following dark acclimation and while kept in the dark, primary leaflets (∼2 cm long) were sampled under a green light, and their petiolets were immediately immersed in pre-labeled single PCR tubes containing either freshly prepared apoplast tracing solution of 0.05% (w/v) Berberine Hemisulfate (Sigma-Aldrich, B3412-10g) in water, or one of the two control solutions: 90mM potassium thiocyanate (KSCN; Sigma-Aldrich, 207799-100G), dissolved in water or water alone. Samples were incubated in the dark for 2 hours, ensuring only the petiolets were completely immersed in the solution throughout the incubation period. Samples incubated with Berberine Hemisulfate were then transferred to new PCR tubes containing KSCN for 2 hours to induce precipitation of berberine thiocyanate within the tissues. Finally, images of the abaxial side of the leaflets were acquired on a Zeiss Discovery V12 fluorescence stereoscope equipped with an AxioCam MR3 camera and an X-Cite 120 illuminator, using a GFP filter set (Ex 470/40, DM 495 LP, Em 525/50).

The tracing of symplastic transport is based on Jiang et al. (Jiang et al., 2019), with modifications described above for the plants’ dark acclimation. Leaflets were sampled (n=6), and their petiolets were immersed in pre-labeled single PCR tubes containing either freshly prepared symplastic tracing solution of 5 μM 5-(and-6)-Carboxyfluorescein Diacetate (CFDA; Sigma-Aldrich, C195) or water. Samples were incubated in the dark for 2.5 hours and imaged as described above. Plasmodesmal permeability was independently quantified following the Drop-ANd-See (DANS) assay (Cui et al., 2015) on the primary leaflet (n=6) from the fourth fully expanded leaf.

#### Image Analysis

Because CoroNa Green is an intensity-based dye that predominantly reports vacuolar Na⁺ in intact plant cells, measurements were interpreted as relative enrichments within specific cell populations rather than as absolute cytosolic concentrations. The intensity of the CoroNa Green signal was quantified using ImageJ version 1.54g. All confocal Z-stack images were projected using maximal intensity projection. Next, for each image, we split the image channels and analyzed only the green channel for area, mean, and integrated Density. Using the freehand tool, the region of interest (ROI) within the leaf cross-section, as well as the background regions of the image (outside the cross-section), was selected. Five to six technical measurements were obtained for the ROI and background regions in each image. Technical measurements were averaged, and the intensity of CoroNa Green signal was calculated based on corrected total cell fluorescence (CTCF) using the following equation: Integrated Density(ROI) - Area(ROI) × Mean(Background); all images were processed with identical thresholds. To estimate the number of Na^+^ accumulating cells among different leaf cell populations, we first defined the cell population of interest within and around the midvein: adaxial parenchyma, abaxial parenchyma, mesophyll, bundle sheath, pericycle, and inner vascular bundle cells. For each maximal intensity projection image, we applied Adjust → Color Threshold with the following fixed settings: Color space = HSB, Threshold color = B&W, Method = Default, Dark background = on. Thresholds for Brightness were set to 55-255 (Hue and Saturation left at full range), producing a binary mask of CoroNa Green-positive pixels. Next, counts were recorded for each cell population of interest and expressed as a numerical value. All images were processed with identical parameters. Quantification of CFDA signal intensity was performed as described in Cui et al. (Cui et al., 2015). Some images were adjusted for brightness and contrast using ImageJ and assembled into figures in PowerPoint.

#### RNA-Sequencing and Data Analysis

M82, *S. pennellii*, and IL6-4 plants were grown as described above. Following 13 days of irrigation with salt water, primary leaflets from the second fully expanded leaf were sampled and immediately flash-frozen in liquid nitrogen. RNA was extracted from four leaf samples (i.e., biological replicates) per genotype and condition using the Direct-zol RNA MiniPrep Plus kit (Neta, RPI-ZR2053). Sequencing libraries were generated as previously described in (Manzano et al., 2025). RNA-seq libraries were pooled and sequenced using the Illumina NovaSeq (PE150). Sequences were pooled, and trimmed using Trim Galore (v.0.6.6). Genome indices were built with STAR (Pebesma & Bivand, 2023) for *S. lycopersicum* SL4.0 (ITAG4.0) (Tomato Genome, 2012) and *S. pennellii* v2.0 (Bolger et al., 2014) (--sjdbGTFtagExonParentTranscript Parent). Trimmed reads were aligned with STAR, and gene counts were generated with HTSeq-count against the respective GFF3 annotations (--type exon, --idattr Parent). To select a reference genome, we compared mapping statistics (e.g., uniquely mapped reads, multimapping, and mismatch rate), annotation completeness/quality, and sample clustering using principal component analysis (PCA) across the three genotypes and conditions. Because mapping metrics were largely similar, PCA structures were nearly indistinguishable between references (**Supplemental Fig. S5A-D**), and the *S. lycope*rsicum annotation is more complete and better curated, we used the *S. lycopersicum*-based alignments for all downstream differential expression analyses. Differentially expressed genes (DEGs) were identified using the limma R package as described in (Manzano et al., 2025). Genes with an adjusted *P* value (P_adj_) ≤ 0.1 were considered differentially expressed (**Supplementary Table S1**). The MapMan annotation file, based on ITAG4.0 (https://mapman.gabipd.org/mapmanstore), was used for functional gene annotations. Since we focused on DEGs within the introgressed region of IL6-4, we used the following approach to identify them: we intersected the 146 *S. pennellii* genes harboring in bin d-6G (IL6-4), based on Spenn-v2-cds-annot.fa file (https://solgenomics.net/ftp/genomes/Solanum_pennellii/), with the 162 *S. lycopersicum* genes that were introgressed out of d-6G, based on Chitwood 2013 and following update to ITAG3.2. The orthology relationship between the two species was determined using an ortholog pair file kindly provided by Prof. Julin Maloof (UCD). BLASTn (https://blast.ncbi.nlm.nih.gov/Blast.cgi) was used to further identify *S. lycopersicum* orthologs among 21 *S. pennellii* genes that were absent from the ortholog pair file. Altogether, *S. lycopersicum* orthologs were assigned to 139 *S. pennellii* genes (**Supplementary Table S2**).

#### RNA Extraction and Real-Time (RT) PCR

To validate the expression of *PDLP1* (Solyc06g083750) in M82 and its orthologs in IL6-4 and tomato wild relatives (*S. pennellii*, *S. pimpinellifolium*, and *S. peruvianum*), plants were grown, and RNA was extracted from primary leaflets from the second fully expanded leaf as described above for the RNA-seq experiment with 3-4 biological replicates per genotype and treatment. First-strand cDNA was synthesized using SuperScript^TM^ III reverse transcriptase (ThermoFisher, 18080-044) following the manufacturer’s instructions. RT-qPCR was performed using PowerUp™ SYBR™ Green Master Mix (ThermoFisher, A25742) on the CFX96TM Real-Time PCR Detection System (Bio-Rad Laboratories Inc.). *PDLP1*-specific primers were designed using the *S. lycopersicum* and *S. pennellii* genomes in Primer-BLAST software (Ye et al., 2012) (**Supplementary Table S4**). The SnapGene software (https://www.snapgene.com/) was used to align the CDS sequences of *PDLP1* orthologs from the four tomato species, ensuring that the amplicon sequences were identical. The 2^−ΔΔCT^ method (Livak & Schmittgen, 2001) was used to normalize and calibrate transcript values relative to the housekeeping gene Expressed (SGN-U346908) (Exposito-Rodriguez et al., 2008).

#### Phylogenetic Analysis

The phylogenetic tree was generated using the method described by Kajala et al. (Kajala et al., 2021).

#### Statistical Analysis and Reproducibility

Statistical analyses were conducted in the R environment (v. 4.3.3), and derived plots were created using ggplot2 (v. 3.5). For multiple comparisons between genotypes, a two-way or one-way ANOVA was performed, followed by a Tukey-Kramer post hoc test or Dunnett’s test when compared with M82, respectively. Enrichment of Na^+^ accumulation within a specific cell population was determined based on Fisher’s exact test. Correlation between Na^+^ quantification using ICP-MS or *PDLP1* expression and fluorescent intensity of CoroNa Green signal, respectively, was done using the Spearman method. Statistical analysis of phenotypic stability within the IL population was done using Student *t*-tests in Microsoft Excel v.1808. Individual *P* values for differences of selected ILs between M82 and S. pennellii under control and salt stress conditions, respectively, are provided in Supplemental Figure S3. For all box plots, the center depicts the median, and the lower and upper box limits depict the 25th and 75th percentiles, respectively. Whiskers represent minima and maxima. Closed dots depict individual samples. In all cases, the number of individual biological samples is denoted as n. Experiments and representative images involving histological staining were repeated at least twice independently, except for the apoplastic transport assay.

## Data Availability

The data that supports the findings of this study are available in the supplementary material of this or raw sequencing files of mRNA sequencing are available at the Short Read Archive of the National Center for Biotechnology Information (https://trace.ncbi.nlm.nih.gov/Traces/sra) under accession number XXXXX.

## Supporting Information

The datasets supporting the results of this article are included within the article and its additional files.

## Funding

This work was supported by funding from NSF-PGRP 2119820 and NSF-IOS 2118017, an HHMI Faculty Scholar fellowship as well as HHMI Investigator funds to S.M.B. L.S.M. was supported by the Va’adia–BARD Postdoctoral Fellowship (BARD FI-570-2018) and the 2022 Women’s Postdoctoral Career Development Award in Science from the Weizmann Institute of Science. L.S.M and S.M.B were supported by BARD IS-5662-24A. D.E.R was supported by the USDA Hatch project CA-D-PLS-2579-CG

## Acknowledgments

We sincerely thank Prof. Neelima Sinha for sharing seeds from the M82 × *S. pennellii* introgression line population, and the TGRC for providing seeds of the wild tomato species. We also thank Dr. Yuh-Ru Julie Lee for valuable advice on confocal microscopy, and Prof. Julin N. Maloof for kindly sharing an orthology relationship file between *S. lycopersicum* and *S. pennellii*. Many thanks to Eduardo Blumwald for providing advice on experimental design and data interpretation. The authors declare no conflicts of interest. The authors confirm that AIGC tools were not used in developing any portion of this manuscript.

## Author Contributions

LSM and SMB designed the research, LSM performed the research, LSM data analysis and collection, LSM, DER and SMB interpreted data, LSM and SMB wrote the manuscript.

## Notes

### Competing Interest Statement

The authors have declared no competing interest.

