## Supplemental Figures for "Cell-specific Na^+^ accumulation is linked to symplastic transport in tomato leaves"

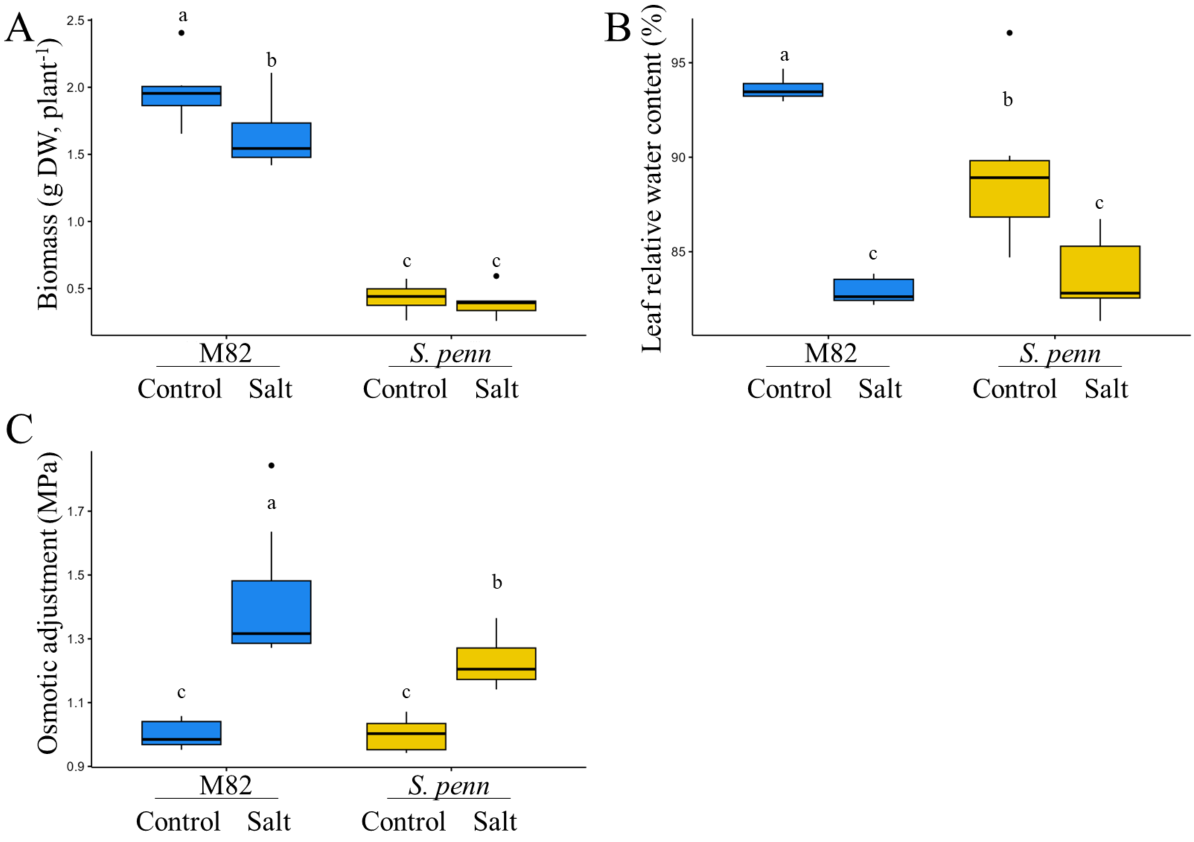


**<u>Figure S1. related to Figure 1:</u> M82 and *S. pennellii* exhibit divergent physiological responses under salt stress conditions**. (**A**) Bar plots of whole plant dry weight (DW, n≥6), (**B**) relative water content of terminal leaflets from the third fully expanded leaf, located below the shoot apex (n≥5), and (**C**) osmotic adjustments of primary leaflets from the second fully expanded leaf (n≥6). Bars are median ± SD. Significant differences (P<0.05), detected by two-way ANOVA followed by Tukey's HSD test, are indicated by different letters.

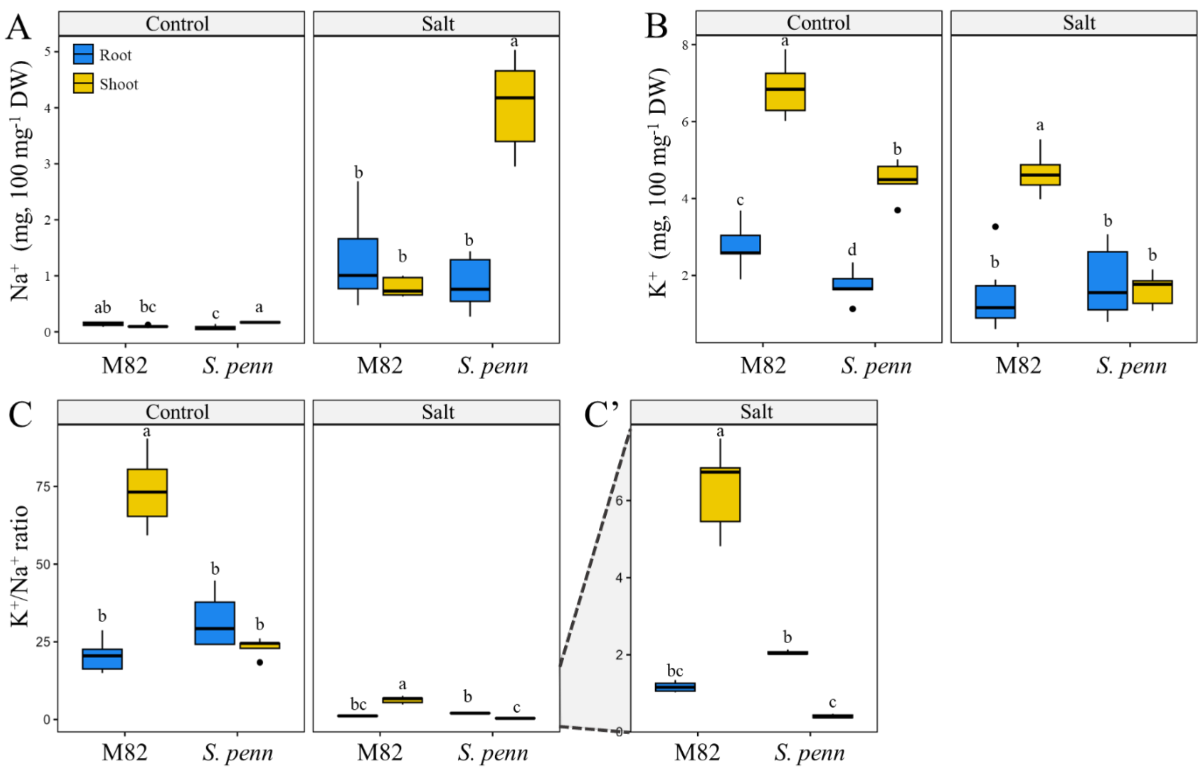


**<u>Figure S2. related to Figure 1:</u> Wild and domesticated tomatoes show different levels of $Na^{+}$ and $K^{+}$ accumulation when exposed to salt stress.** Bar plots of leaf (**A**) $Na^{+}$ and (**B**) $K^{+}$ content and (**C**) $K^{+}/Na^{+}$ ratio (n=6) measured by inductively coupled plasma mass spectrometry. (**C'**) Zoom in on the $K^{+}/Na^{+}$ ratio under salt stress conditions. Bars are median ± SD. Significant differences ($P<0.05$) within each condition, detected by two-way ANOVA followed by Tukey's HSD test, are indicated by different letters.

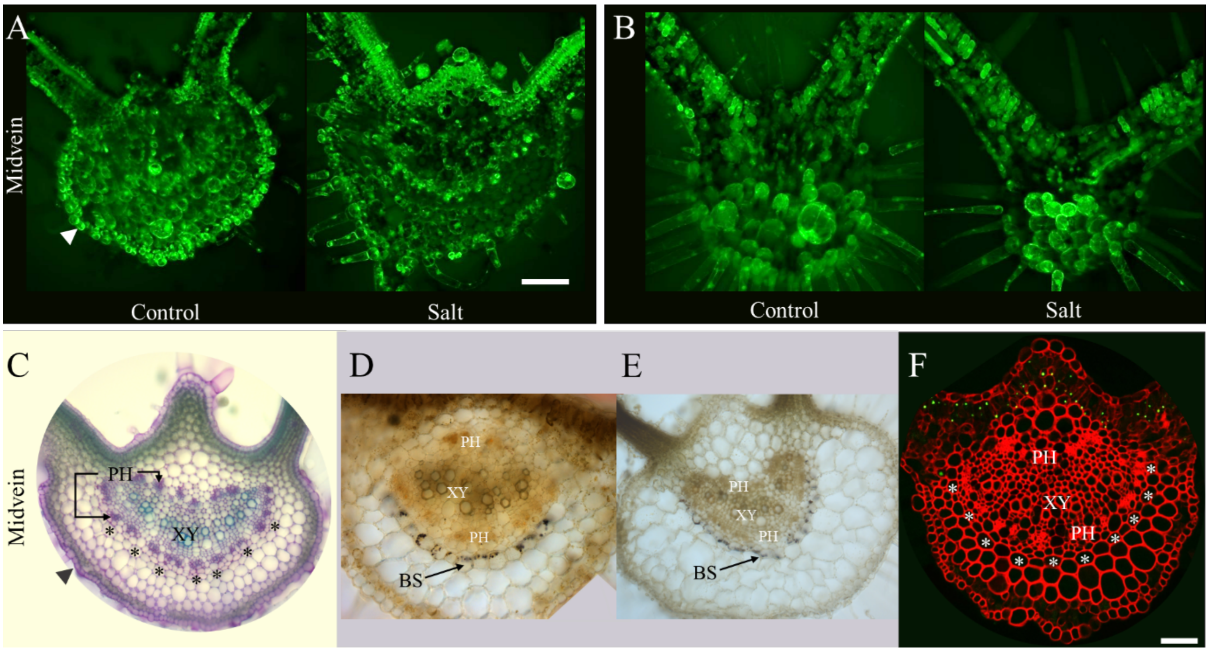


**<u>Figure S3, related to Figure 1:</u>** Histological analysis of leaf samples from M82 and *S. pennellii*. Representative live cross sections of the leaf midvein from five-week-old plants stained with fluorescein diacetate (FDA) to confirm cell viability. Cells in the cross sections of (**A**) M82 and (**B**) *S. pennellii* from both growth conditions are viable, as indicated by a strong fluorescent signal reflecting cell integrity and activity. Scale bar represents 110 µm. To identify tomato leaf cell types, especially the bundle sheath, cross sections were stained with (**C**) toluidine blue, a metachromatic dye used to differentiate primary and secondary cell walls (e.g., xylem [XY] and phloem [PH] cells), iodine-potassium iodide (IKI), a starch-specific dye that stains amyloplasts within the bundle sheath [BS] (**D**) of M82 and (**E**) *S. pennellii* , and (**F**) Congo red, which stains cellulose to visualize different cell layers. The bundle sheath is indicated by asterisks in panels **C** and **F**. The scale bar indicates 100 µm.

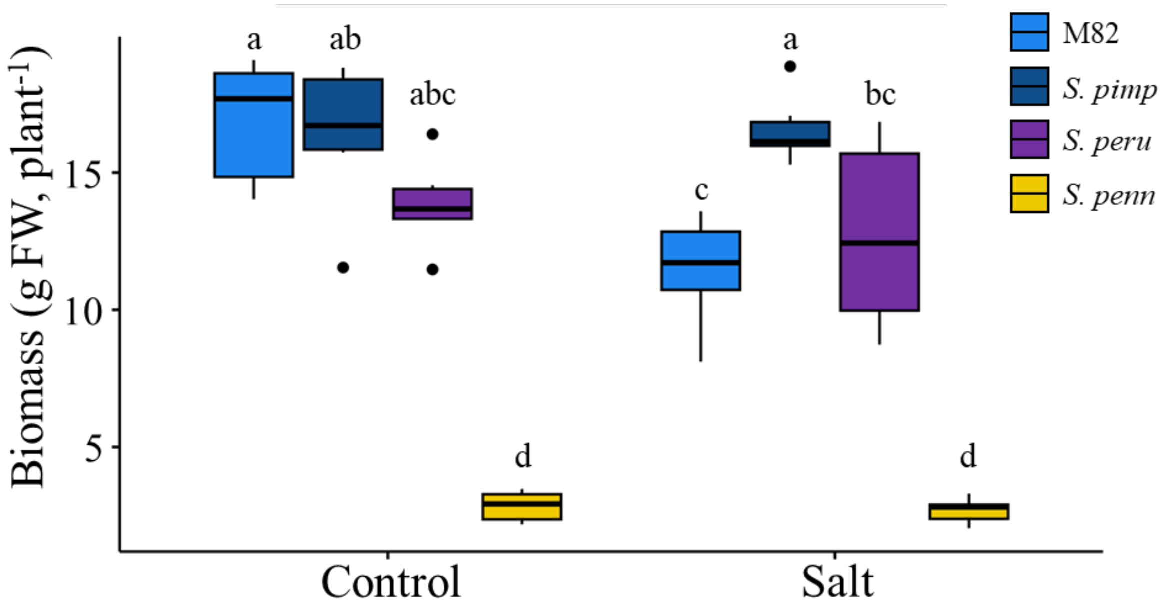


**<u>Supplemental Figure S4. related to Figure 2:</u> Biomass accumulation of salt-tolerant wild relatives of tomato.** Box plots of fresh weight (FW) of tomato wild relatives grown under control and salt stress conditions. Bars are median ± SD (n=6). Significant differences ($p$<0.05), detected by two-way ANOVA followed by Tukey's HSD test, are indicated by different letters.

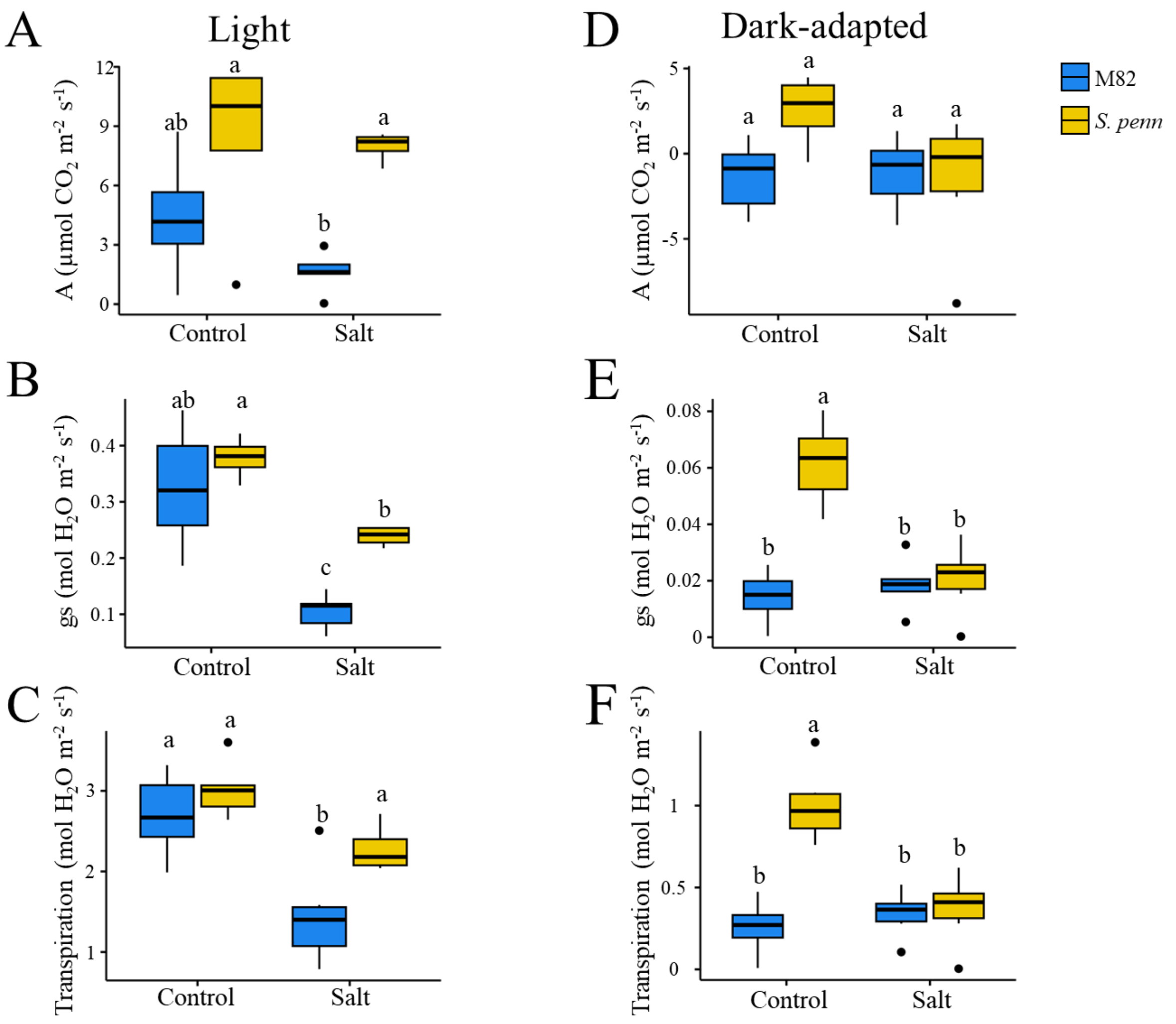


**Supplemental Figure S5. related to Figure 3: Photosynthesis in M82 and *S. pennellii*.** Photosynthesis rate (A) (**A & D**), stomatal conductance (gs) (**B & E**), and transpiration rate (**C & F**) under light conditions and after two hours of dark adaptation in control and salt-stressed tomatoes. Measurements were taken on the abaxial surface of the terminal leaflet from the third or fourth fully expanded leaf. Bars show median ± SD (n≥5). Significant differences ($P<0.05$), determined by two-way ANOVA followed by Tukey's HSD test, are indicated by different letters.

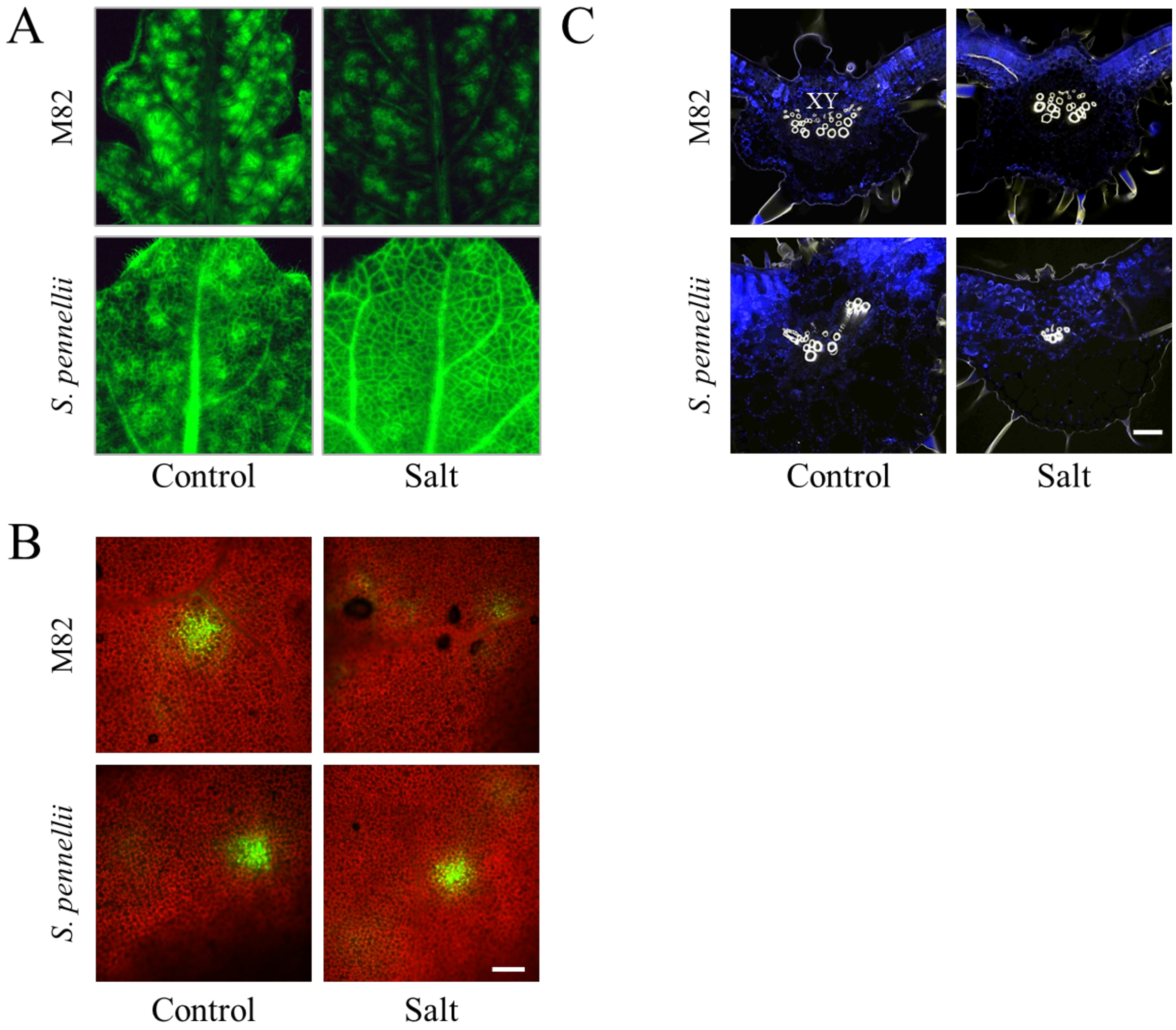


<u>**Supplemental Figure S6. related to Figure 3:**</u> **Qualitative data on symplastic transport and staining for an apoplastic barrier in tomato leaves**. (**A**) Representative fluorescent images of leaflets subjected to 5(6)-Carboxyfluorescein diacetate (CFDA) petiole feeding (indicated in green). M82 and *S. pennellii* were grown under control and salt stress conditions and subjected to two hours of dark adaptation before being treated with the symplastic tracer. (**B**) Representative confocal images of CFDA absorption (green) at the adaxial surface of the leaflet. Chloroplast autofluorescence is observed as a red background. Images indicate symplastic movement of CF from the adaxial to the abaxial epidermis. Scale bar = 200 µm. (**C**) Representative confocal images of midvein cross-sections stained with Auramine O dye (yellow) from M82 and *S. pennellii* under salt stress and control conditions. Chloroplast autofluorescence is visible in blue. The xylem cell wall and the epidermal cuticle are stained with Auramine O. There is no evidence of deposition of hydrophobic polymers such as lignin, suberin, and cutin around the midveins in either condition in either species. Scale bar = 110 µm.

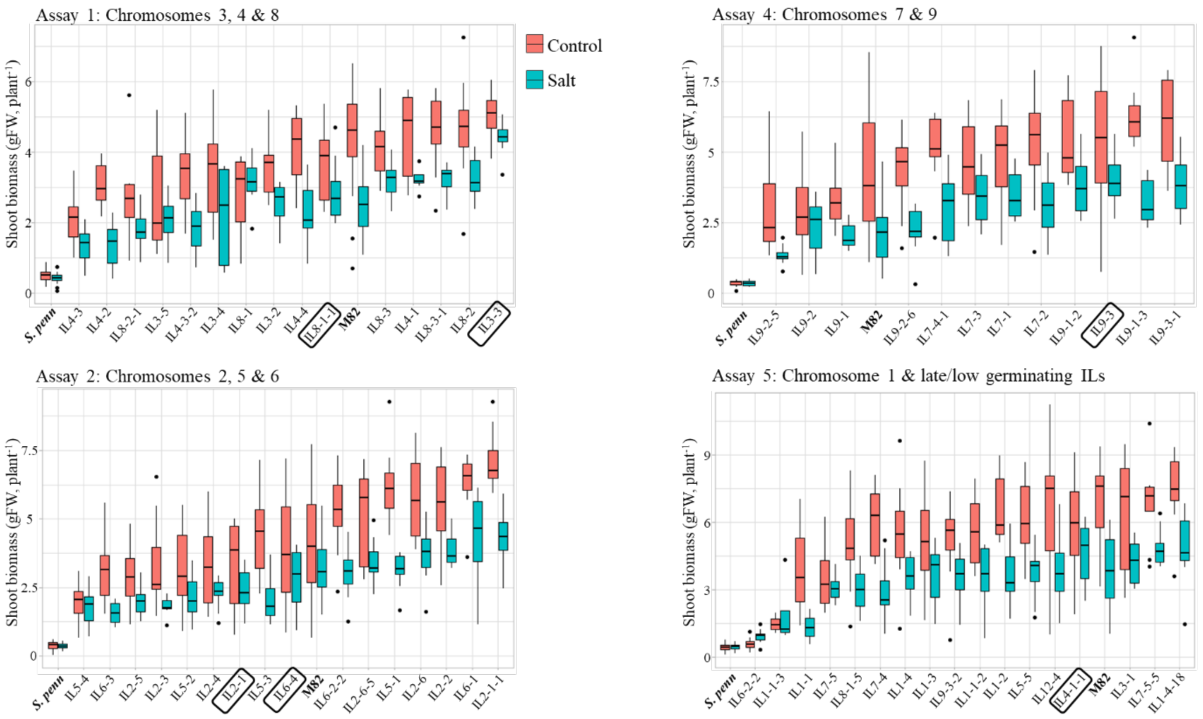


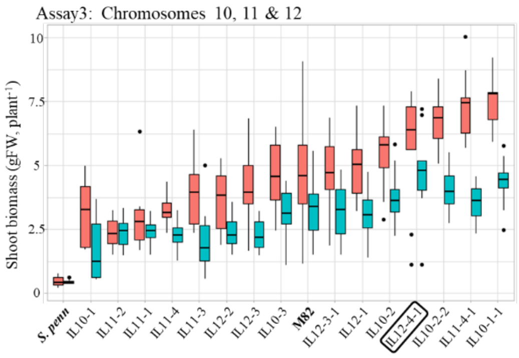


| Assay | Genotype | Mean IL Salt | Mean IL Ctrl | Δ IL Salt-Ctrl | pV (S-C) | Mean M82 Ctrl | Δ Ctrl IL-M82 | pV (IL-M82) |
|---|---|---|---|---|---|---|---|---|
| 1 | IL3-3 | 4.41 | 5.05 | -0.6 | 0.01 | 4.54 | 0.51 | 0.23 |
| 1 | IL8-1 | 3.17 | 3.81 | -0.6 | 0.21 | 4.54 | -0.73 | 0.26 |
| 3 | IL12-4-1 | 4.74 | 5.65 | -0.9 | 0.19 | 4.63 | 1.02 | 0.18 |
| 2 | IL2-1 | 2.37 | 3.31 | -0.9 | 0.06 | 4.19 | -0.88 | 0.22 |
| 2 | IL6-4 | 2.81 | 3.9 | -1.1 | 0.08 | 4.19 | -0.29 | 0.70 |
| 5 | IL4-1-1 | 4.57 | 5.67 | -1.1 | 0.10 | 6.91 | -1.24 | 0.12 |
| 4 | IL9-3 | 4.05 | 5.35 | -1.3 | 0.07 | 4.15 | 1.2 | 0.14 |

**<u>Supplemental Figure S7. related to Figure 4:</u> Screening of introgression line population for phenotypically stable lines.** Box plots show the fresh weight of the IL population under both control and salt stress conditions. Each graph represents an independent assay that includes the wild (*S. pennellii*, n≥16) and domesticated (M82, n≥20) parents, as well as ILs of specific chromosomes (n≥10) for each treatment. The table lists the seven selected ILs that demonstrate phenotypic stability under salt stress, meaning they exhibit a non-significant or minor reduction in fresh weight under salt stress conditions, similar to *S. pennellii*, and show no significant difference from M82 under control conditions. Significant differences (P<0.05) were detected using the Student t-test. Control conditions are indicated in red, and salt stress conditions in turquoise.

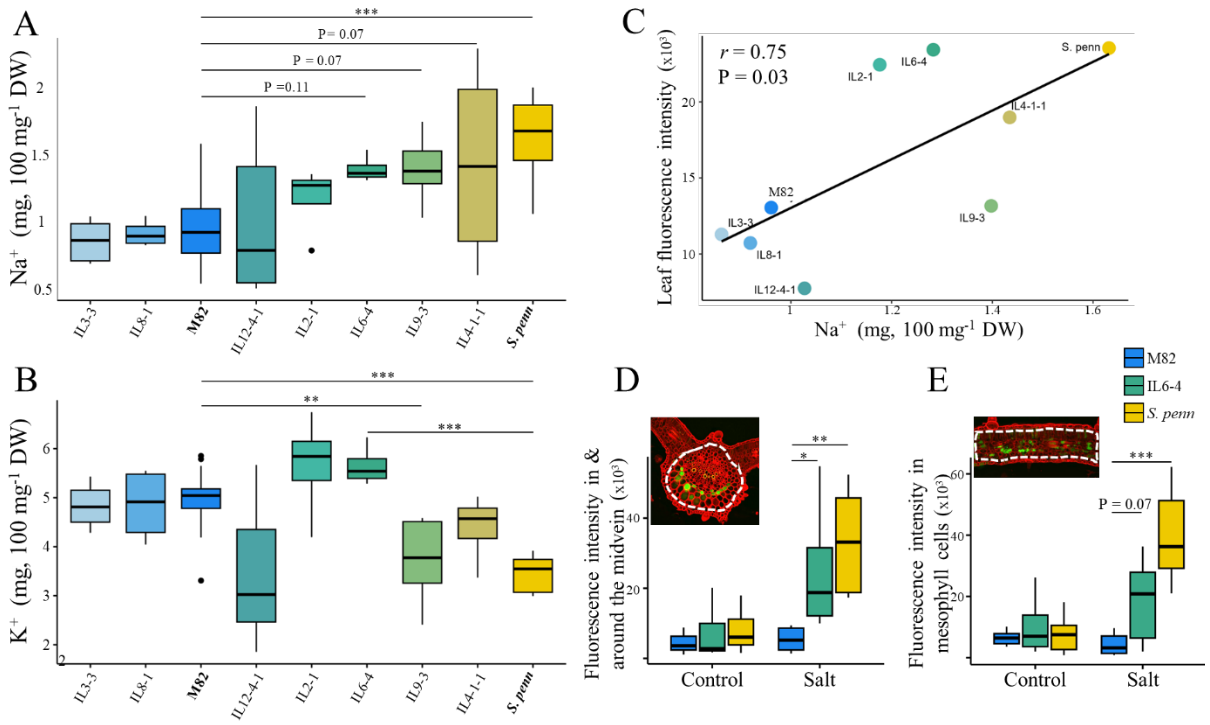


**<u>Supplemental Figure S8. related to Figure 4:</u> Quantification of ion content in leaves of selected introgression lines.** Box plots of (**A**) $Na^+$ and (**B**) $K^+$ content in leaves of the seven selected ILs grown under salt stress conditions. Dry weight, DW. Bars represent median ± SD (n≥4). Differences were tested with one-way ANOVA followed by a one-sided Dunnett's test comparing each genotype to M82 (**A**) and M82 or *S. pennellii* (**B**). Asterisks indicate significance levels (**P≤ 0.01, ***P ≤ 0.001) between M82*, S. pennellii*, IL6-4, and IL9-3. *The same data of M82, S. pennellii, and IL6-4 are shown in Fig. 2A.* (**C**) Correlation between $Na^+$ content measured with the $Na^+$ specific dye CoroNa green and ICP-MS across the selected ILs, M82, and *S. pennellii*. Box plots show $Na^+$ accumulation, measured as corrected total cell fluorescence (CTCF) (**D**) in and around the midvein and (**E**) in the blade mesophyll (n≥5). Quantification relies on the CoroNa Green signal. White dashed lines mark the areas selected for signal measurement. Significant differences identified by one-way ANOVA followed by Dunnett's test are indicated with asterisks (** P≤0.01, and *** P≤0.001).

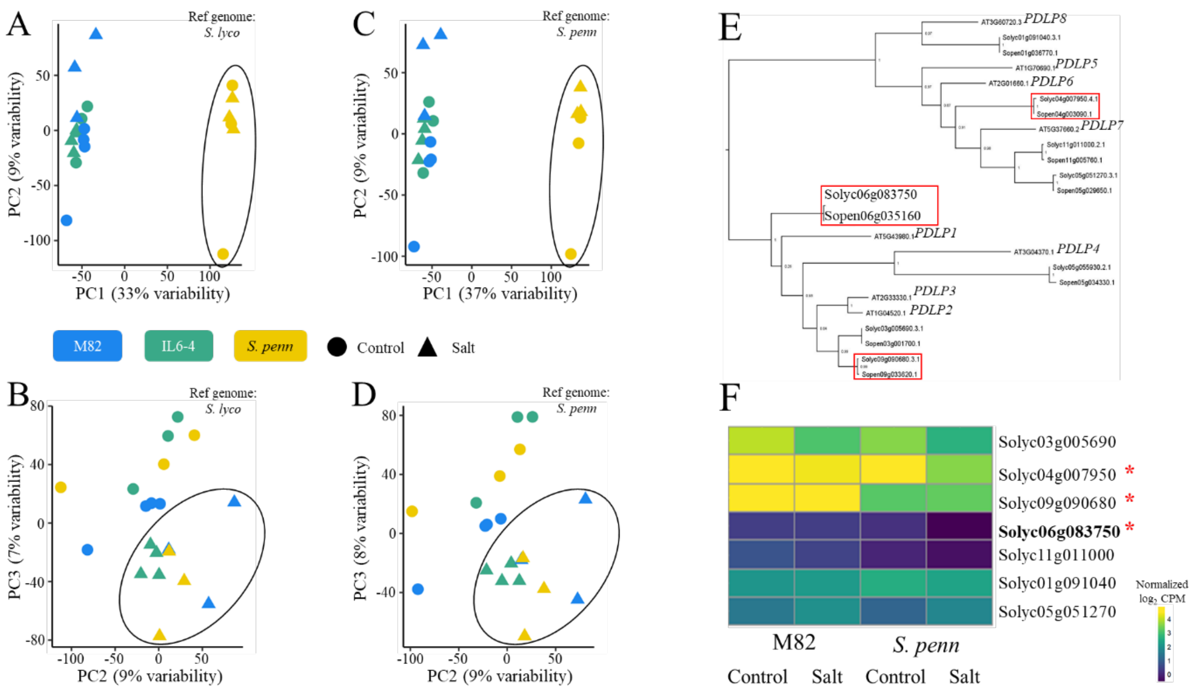


**<u>Supplemental Figure S9. related to Figure 4:</u> Analysis of bulk RNAseq data.** Plots of principal component (PC) analysis of normalized log2 expression values mapped to *S. lycopersicum* (**A** & **B**) or *S. pennellii* genome (**C** & **D**) show similar sample clustering (n≥3). (**E**) Phylogenetic tree of genes encoding *plasmodesmata-located proteins* (*PDLPs*) in Arabidopsis, M82*,* and *S. pennellii*. Differentially expressed *PDLPs* between the two tomato species under salt stress conditions are indicated by red rectangles. (**F**) Heatmap showing expression levels of *PDLP* genes in M82 and *S. pennellii* under control and salt stress conditions. *PDLP1* (Solyc06g083750) and two additional *PDLPs*, marked with red asterisks, show higher expression in M82 compared to *S. pennellii* under salt stress conditions (FDR≤0.1).
